# A fast numerical integration scheme for clonal expansion processes on graphs

**DOI:** 10.1101/2025.01.13.632690

**Authors:** Chay Paterson, Miaomiao Gao, Joshua Hellier, Georg Luebeck, David Wedge, Ivana Bozic

## Abstract

Compound birth-death processes are widely used to model the age-incidence curves of many cancers [1]. There are efficient schemes for directly computing the relevant probability distributions in the context of linear multi-stage clonal expansion (MSCE) models [2]. However, these schemes have not been generalised to models on arbitrary graphs, forcing the use of either full stochastic simulations or mean-field approximations, which can become inaccurate at late times or old ages [3, 4]. Here, we present a numerical integration scheme for directly computing survival probabilities of a first-order birth-death process on an arbitrary directed graph, without the use of stochastic simulations. As a concrete application, we show that this new numerical method can be used to infer the parameters of an example graphical model from simulated data.

## 1 Introduction

The relationship between age and cancer incidence has provided crucial biological insights since it was first noted by Armitage and Doll [5, 6, 7]. It was recognised that the incidence could be modelled as the hazard function of a compound random process, with multiple successive steps corresponding to accumulating mutations in a population of pre-neoplastic cells [5, 8]. This class of mathematical models established a paradigm for studies of cancer, age, and public health; resulting in the discovery of tumour suppressor genes, and allowing detailed, cohort-specific models of environmental risk factors [2, 7, 9].

The most sophisticated models incorporate selection and cell turnover in detail, and allow inferences about the likely number of “hits” involved in the initiation of a given type of cancer to be drawn from data on patient age at diagnosis [1, 10]. Historically, patient age, cohort (via date of birth), and family history were the only observables considered. Modern sequencing technologies can derive much more detailed information about what driver mutations initiated a given tumour, and incorporating this information into multi-stage clonal expansion (MSCE) models is one of the goals of this research. These models include four types of process:

- Mutation (by asymmetric division): *j* → *j* + *k*, at rate *µ*_*jk*_*N*_*j*_
- Birth (by symmetric division): *j* → *j* + *j*, at rate *α*_*j*_*N*_*j*_
- Death: *j* → ∅, at rate *β*_*j*_*N*_*j*_
- Immigration (to approximate mutations appearing in a large unobservable population [1]): ∅ → *j*, at rate *κ*_*j*_

labelling different cell types *j* and *k*, and the numbers of these types *N*_*j*_ and *N*_*k*_, respectively. In most existing MSCE models, the different cell types *j* and *k* represent stages in a linear progression, such as the number of driver mutations, and are indexed with integers [1, 10]. In more recent work, the types *j* are considered as nodes on a directed graph, in order to distinguish between different mutational mechanisms, such as point mutations and copy number alterations [3, 4].

There are thus four possible types of event for each subpopulation *N*_*j*_: mutation, birth, and death obey first-order kinetics, and immigration obeys zeroth-order kinetics. The quantity of the most interest is the probability *S*_*f*_ (*t*) that no primary tumour of type *f* has appeared by a given age *t*. From *S*_*f*_ (*t*), one can define the hazard function *h*_*f*_ (*t*),

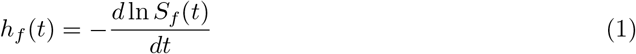

which models the age-related incidence [1].

Unlike the linear case, in which we can identify the probability to have no cancerous cells *P*(*t, N*_*f*_ = 0) with the survival curve *S*_*f*_ (*t*), when there are multiple end states the proper definition of *S*_*f*_ is more subtle. A detailed probabilistic treatment of this subtle technical point is given in appendix A. These multiple end states represent different sets of clonal alterations in the initiated primary tumour: these mutations will be present in every descendant cell in the resulting tumour. As stochastic processes, probabilities (and thus hazard functions) for these models can be computed in a natural way by sampling many replicates of stochastic simulations using e.g. Gillespie’s algorithm. However, stochastic simulations are computationally expensive. Methods to directly estimate *S*_*f*_ (*t*) and *h*_*f*_ (*t*) numerically, given some set of model parameters, are therefore highly desirable for statistical fitting.

In fact, for linear graphs, this type of model is equivalent to *n*-step MSCE models, which have very well-studied methods for reducing the computation of *S*_*f*_ (*t*) to numerical quadrature [1, 2, 10, 11, 12]. These methods study *S*_*f*_ (*t*) via an intermediate generating function, conventionally denoted Ψ or Φ, and use Kolmogorov backward equations to derive recursive formulae for the probabilities *S*_*f*_ (*t*) [11, 8]. The probabilities *S*_*f*_ (*t*) may then be evaluated using an numerical integration scheme such as Gaussian quadrature [1]. In contrast to Kolmogorov backward equations, Kolmogorov *forward* equations have been less intensively studied in the context of MSCE models [10, 13]. We are also aware of recent proposals by Tang et al. that extend similar methods based on Kolmogorov backward equations to describe clonal expansion processes on graphs [14].

The mutational transitions studied in MSCE models of cancer incidence have been mainly limited to linear or path graphs; and in the vast majority of cases, including our current work, the kinetics are first order in the precursor cell populations [1, 15, 16]. More general models involving birth-death-mutation processes on arbitrary directed graphs have been studied using stochastic simulations, or approximate solutions derived from mean-field or moment closure approximations [3, 17, 18]. A limited number of analytical results for first-order chemical reaction networks are also known. Notably, monomolecular systems in which there is no more than one product per reaction, are known to be exactly solvable [19, 20, 21, 22, 23].

Due to the importance of stochasticity in biological systems, the discovery of exact analytical, semi-analytical, and numerical solutions is still an important open problem, given that stochastic simulation is computationally expensive [20, 24]. Here, we describe an improved numerical scheme based on Kolmogorov forward equations and the method of characteristics. This algorithm is suitable for studying clonal expansion models with constant coefficients on arbitrary directed graphs [11, 13].

## 2 Methods for estimating *S*_*f*_ (*t*)

The problem being considered is the estimation of the diagnosis-free survival curve *S*_*f*_ (*t*) for a clonal expansion model on a graph *G*. The graph *G* consists of a set of vertices *V*, interpreted as different mutant genotypes, and a set of edges *E*, interpreted as different mechanisms by which driver mutations can be acquired.

Different vertices in *V* correspond to subpopulations with different combinations of genetic alterations. Each subpopulation *N*_*j*_, *j* ∈ *V*, is a random variable that is a non-negative integer, so that *N*_*j*_ ∈ Z_≥0_. The initial value of *N*_*j*_ at age *t* = 0 will be denoted *Z*_*j*_.

There is also a subset of nodes *F* ∈ *V* which are “final” subpopulations. These are interpreted as neoplastic or cancerous cells. Our overarching goal is to compute the probability *S*_*f*_ (*t*) that an individual has survived to age *t* with no primary tumours of type *f*. This is related to the probability that a chosen final node *f* ∈ *F* is unoccupied by time *t*: *P*(*t, N*_*f*_ = 0) (see A).

In general, the rate coefficients {*α*_*j*_, *β*_*j*_, *µ*_*jk*_, *κ*_*j*_} may vary with respect to age or time. However, in this work, the special case where they are constant will be given special consideration, as some dramatic optimisations are then possible. Since the mutation rate is unlikely to decrease with age, we may consider the resulting estimates of *S*_*f*_ to represent an upper bound.

The edge set *E* corresponds to pairs of nodes for which the mutation rate *µ*_*jk*_ *>* 0, and are interpreted as different mutational mechanisms. These may be mutations on different loci, or different biological processes such as deletions, loss of heterozygosity, copy number alterations and so on. For example, the graph may be a tree, with different vertices representing different ancestral genotypes. There are many imaginable applications: a directed graph of this type can encode any string [25].

The set of nodes *V* and edge set *E* together define a graph *G* = (*V, E*) [26]. In contrast to Luebeck and Moolgavkar, we will not assume a specific linear structure for this graph, and will instead study methods suitable for arbitrary graphs *G* [10].

These dynamics imply the following form for the Kolmogorov forward equations:

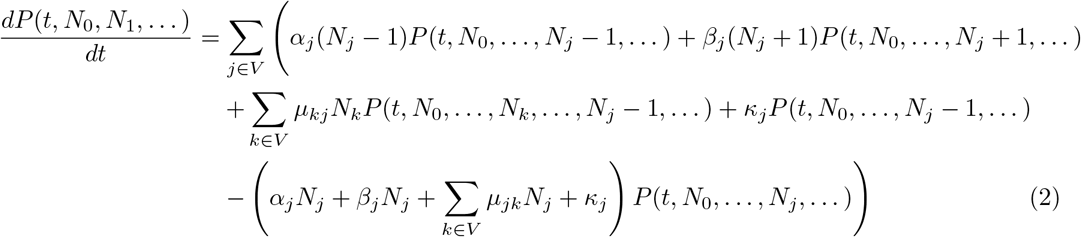

The Kolmogorov forward equations are also known as the chemical master equation [24, 27, 28]. In general, this system of ordinary differential equations is infinite-dimensional, and known solution methods are extremely limited [20]. The only general method for computing arbitrarily precise solutions for the distribution 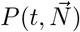 is Gillespie’s stochastic simulation algorithm [29].

We will now briefly review Gillespie’s algorithm, before describing the formal solution of equation (2) via the method of characteristics, and then describe two numerical schemes based on it. The second scheme, described in 2.4, is novel and fast.

### 2.1 Gillespie’s stochastic simulation algorithm

Gillespie’s stochastic simulation algorithm provides a “brute force” method for estimating the distribution *P*(*t, N*_0_, …) in (2), and is widely used to simulate biological processes and chemical reaction networks. The probability for each reaction to fire in a given time interval depends on the populations of different chemical species and the rate coefficients of different reaction pathways. The most important property of the Gillespie algorithm is that it provides an exact solution of the master equation (2) that describes the dynamics of the system, in the sense that the trajectories the algorithm produces are random samples of the probability distribution that solves the master equation (2) [29, 30].

We define the state of the populations in the system to be 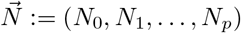 with each *N*_*i*_ the population of species *i*; we index the reaction pathways with integers *j* in 0 ≤*j* ≤*k*; and let the rate of each reaction pathway *j* be 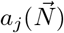. Gillespie’s stochastic simulation algorithm can then be described as follows:

1. Initialisation: set the time *t* = 0, set the populations 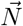 to their initial values 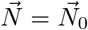, and initialise the random number generator.
2. With the system in state 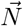, evaluate every 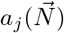 and their sum 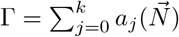.
3. To decide which reaction *r* will happen, generate a uniform random number *X* ∈ [0, Γ]. The reaction *r* is the smallest integer that satisfies

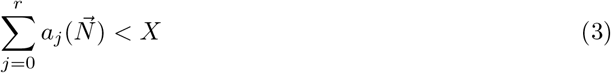 This method of choosing *r* guarantees that the probability that reaction channel *r* is chosen this step is equal to 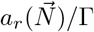.
4. Compute the time *τ* until the next reaction by generating a uniform random number *Y* ∈ [0, 1], and computing

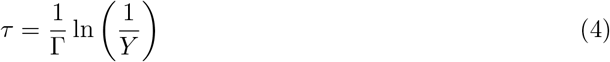
5. Update the state 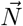 according to the reaction *r*, and increment *t* by *τ*.
6. Iteration: return to step 2 until the termination condition has been met. Example conditions are that the time *t* reaches some maximum value *t*_*max*_, or a specific population *N*_*i*_ reaches a maximum value.

In each iteration, only two calls to the random number generator are required.

### 2.2 Wave equation and formal solution by the method of characteristics

First, we define the generating function Ψ to be the expected value of 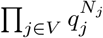 given a vector of conjugate coordinates 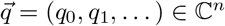 [31].

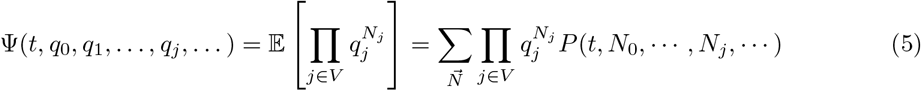

where the sum runs over all possible values of the populations 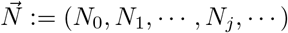. That is, 0 ≤ *N*_*j*_ *<* ∞, for all *j* ∈ *V*. This is essentially the Fourier transform of the probability distribution *P*(*t, N*_0_, …) with respect to the variables *N*_*j*_ [31]. The variables *q*_*j*_ are conjugate to the populations *N*_*j*_, forming Fourier duals in a similar sense to frequency and time.

The probability *P*(*t, N*_*f*_ = 0, *N*_*f*_*′* = 0) that the populations of two final nodes *f* and *f*^*′*^ are both zero at age *t* can then be expressed in terms of Ψ:

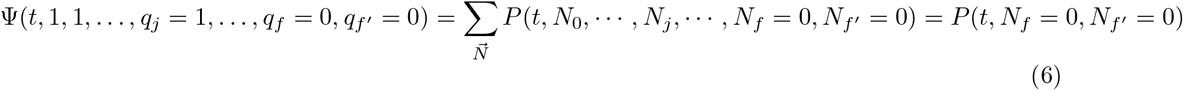

i.e. set *q*_*f*_ = 0 for every node *f* that is unoccupied, and *q*_*j*_ = 1 for every other node. More tersely, *q*_*j*_ = 1 − *δ*_*jf*_ − *δ*_*jf*_*′*, with *δ*_*ij*_ the Kronecker delta. In general, the probability that all nodes *f*^*′*^ in a subset *X* are unoccupied at age *t* can be expressed as:

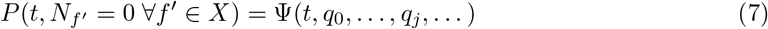

where *q*_*j*_ = 0 if *j* ∈ *X*, and *q*_*j*_ = 1 otherwise. This allows us to compute the diagnosis-free survival curve *S*_*f*_:

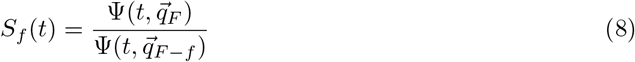

where the coordinate vectors 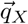 are shorthand for:

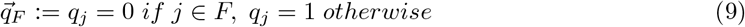

and

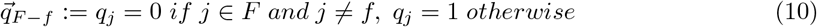

respectively. See A for details of the definition of *S*_*f*_.

When recast in terms of the generating function Ψ, equation (2) becomes the partial differential equation

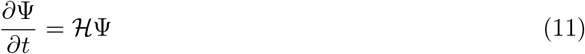

where the hyperbolic differential operator ℋ (or “Hamiltonian”) is

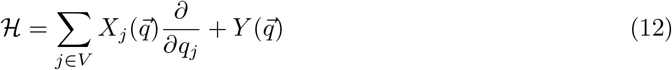

where

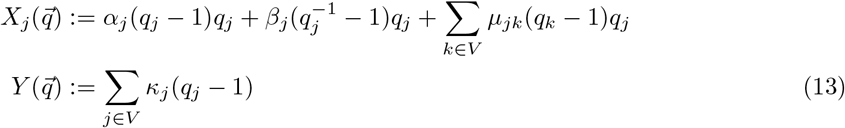

The resemblance between equation (11) and others in mathematical physics is not a coincidence, and has been discussed by other authors [32].

#### Theorem

A general solution to (11) is given by

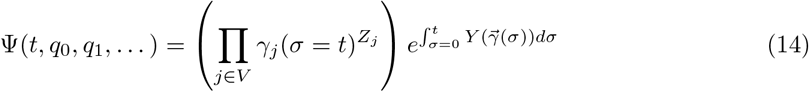

where the characteristics *γ*_*j*_ are solutions of the initial value problem

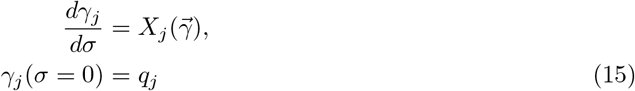

and the vector field 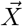 and scalar field *Y* are defined to be

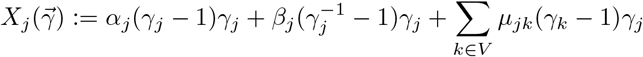

and

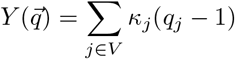

and 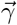 is shorthand for (*γ*_0_, *γ*_1_,· · ·, *γ*_*j*_,· · ·).

Values of Ψ can thus be computed for any given set of initial conditions 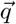. As an immediate corollary, this allows *S*_*f*_ (*t*) to be calculated: the relevant initial conditions are given by (6), with *q*_*j*_ = 1 for all *j* ≠ *f*, and *q*_*f*_ = 0. Hence,

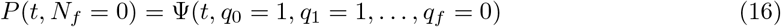

#### Proof

The wave equation (11) can be placed into a conservation form along a vector field 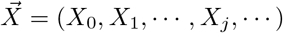:

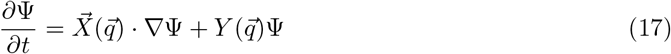

This can be solved using the method of characteristics [33]. Briefly, we seek a parametrised family of curves *τ*(*σ*), 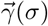 for which

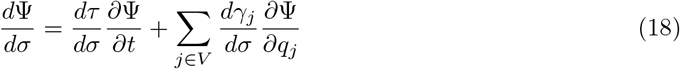

and *τ*(0) = *t*, 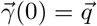. An appropriate family of curves is

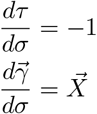

Clearly *τ*(*σ*) = *t* − *σ*. The characteristics 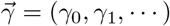 will have a more complicated form. Along these curves, equation (17) takes the form

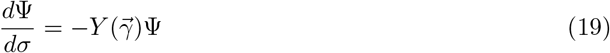

where

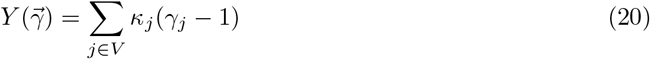

Integrating (19) and rearranging for 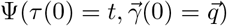 yields

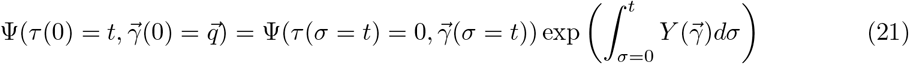

Given that the initial conditions are known,

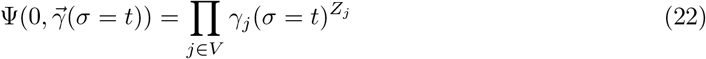

substituting this results in the formal solution (14).

### 2.3 Numerical integration schemes based on Kolmogorov forward equations

The formal solution (14) via the method of characteristics opens up many possibilities for numerical methods [27]. In principle, any numerical integration scheme that is appropriate for systems of ODEs can be used to solve (14), and thus give a numerical approximation for *S*_*k*_(*t*). As these methods can be expected to be “fast” in comparison to sampling with Gillespie’s algorithm, and are based on Kolmogorov forward equations instead of Kolmogorov backward equations (like the known methods from [1, 8, 10]), we will refer to these as “fast forward” methods.

#### Non-constant parameters: Quinn’s method

When the coefficients {*α*_*k*_(*t*), *β*_*k*_(*t*), *κ*_*k*_(*t*), *µ*_*jk*_(*t*)} have an explicit time dependence, the following scheme by Quinn can be used [13]. Explicit time dependence is relevant to e.g. tobacco smoking and possibly inflammatory conditions linked to pre-cancer such as Barrett’s oesophagus. In the following, the coefficients are assumed to have a specified functional form, and e.g. *α*_*j*_(*t*) represents a function call.

For each value of *t*, new characteristic curves *γ*_*j*_(*σ*) (with initial conditions *q*_*j*_) and weights 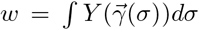 are computed in a “two pass” procedure, using Euler integration to update the *γ*_*j*_, and the rectangle rule to update the weights. These characteristic curves can then be substituted into (14), and corresponding values for *P*(*N*_*j*_ = 0) computed.

1. Choose and fix a time step Δ*t*.
2. For each time of interest *t*…
3. Initialise *σ* = 0, *w* = 0 and *γ*_*j*_ = *q*_*j*_ for all *j*.
4. While *σ < t*:
5. Update *γ*_*j*_ = *γ*_*j*_ + *X*_*j*_Δ*t* for all *j*. (Euler integration)
6. Update 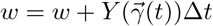.
7. Update *σ* = *σ* + Δ*t*.
8. Store the values 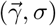.
9. Using the cached values of 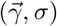 compute 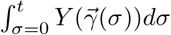 and hence 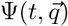.

For any explicit time-stepping procedure with step size Δ*t*, the number of operations to compute one of these characteristic curves will be *N* = ⌈*t*_*max*_*/*Δ*t*⌉. To compute values of *P*(*N*_*j*_ = 0) at all *N* ages in between *t* = 0 and *t* = *t*_*max*_ will therefore require 𝒪(Δ*t*^−2^) operations. This is explicitly described in Crump’s review and Quinn’s original paper [11, 13].

### 2.4 Constant coefficients: a one-pass method

Inspecting the formal solution (14) of the Kolmogorov forward equations immediately suggests other numerical schemes along the lines of Quinn’s. We will describe a simple implementation based on *improved* Euler integration: more sophisticated schemes are obviously possible. However, when the coefficients {*α*_*k*_, *β*_*k*_, *κ*_*k*_, *µ*_*jk*_} are independent of time, a more radical optimisation of Quinn’s method is possible.

Consider what happens to (14) when *t* is replaced by *t* + Δ*t*, where Δ*t* is a finite time step. Equation (14) is valid at all times, so it must be true that

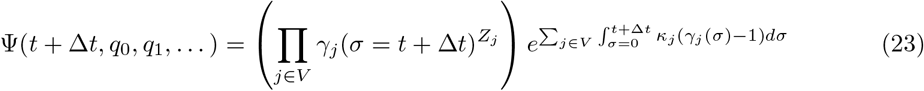

Each characteristic solves (15), so

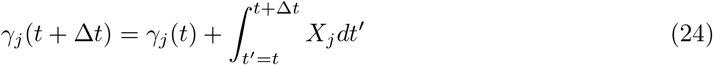

which can be computed with an appropriate numerical integration procedure, for example improved Euler integration:

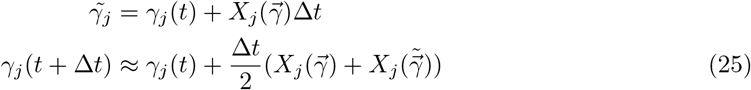

(or a relative such as a higher-order Runge-Kutta, etc.).

The remaining term can also be approximated numerically. For brevity, call

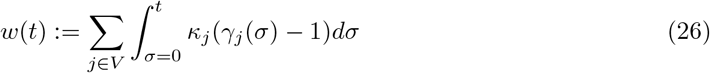

and notice that

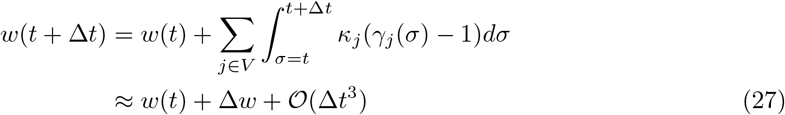

where the latter approximation integrates *w*(*t*+Δ*t*) using improved Euler integration. The errors in this approximation must be of the same order as the errors in the integration of the characteristics 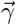 in equation (25). Equation (23) can now be written

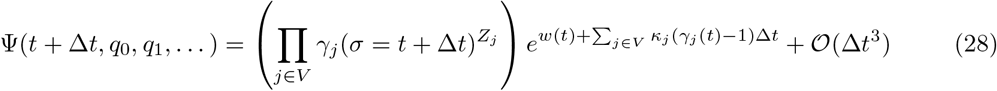

Therefore, to calculate the value of Ψ at time *t* + Δ*t*, we only need to know what the values of 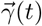 and *w*(*t*) were. This shows that the generating function 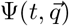 can be computed numerically, by applying a time-stepping procedure to the characteristics 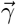 and the weight *w*(*t*). A concrete implementation in pseudocode might be:

1. Choose and fix a time step Δ*t* and maximum time *t*_*max*_.
2. Initialise *t* = 0, *w* = 0 and *γ*_*j*_ = *q*_*j*_ for all *j*.
3. Update *γ*_*j*_ for all *j* with an improved Euler step.
4. Update 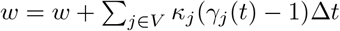.
5. Update *t* = *t* + Δ*t*.
6. Compute 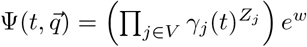.
7. If *t* is less than *t*_*max*_, go to 3. Otherwise, exit.

The only dynamical quantities that need to be stored and updated are *t*, the weight *w*, and the characteristics *γ*_*j*_. A new value of 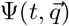 may be computed and returned at every time step. Only one set of characteristics need to be computed between *σ* = 0 and *σ* = *t* to get a complete set of values for Ψ. This scheme terminates after *N* = ⌈*t*_*max*_*/*Δ*t*⌉ ∼ Δ*t*^−1^ iterations. This is asymptotically much faster than Quinn’s method – requiring 𝒪(*N*) operations instead of 𝒪(*N*^2^). Because this numerical method is asymptotically faster than both stochastic sampling with Gillespie’s algorithm and Quinn’s proposal, and based on Kolmogorov forward equations instead of the more Kolmogorov backward equations, we will refer to it as a “fast forward” method.

This can be seen as a prototypical example of a more general class of such methods, because more sophisticated schemes are easy to imagine. For example, improved Euler integration of the characteristics *γ*_*j*_ could be replaced by an adaptive time step or one of a family of higher-order Runge-Kutta methods. Two-pass methods may be worth revisiting to allow for time-dependent coefficients. Many alternatives and optimisations may be possible, but we leave these to future work in order to focus on an example application.

### 2.5 Application to statistical inference: maximum likelihood optimisation

The fast and efficient computation of survival curves *S*_*f*_ (*t*) suggests an application to statistical inference: namely, can this be used to estimate parameters from datasets? This would allow the parameters of graphical MSCE models to be fitted to data in a way that was not practical using random sampling [16, 14, 4, 3]. In this section, we propose an algorithm to achieve this using a simulated dataset that is based on maximum likelihood estimation [2], and study its performance.

Our one-pass fast forward method can be embedded in more complex workflows in which the Gillespie algorithm would be too slow to be practical. As an example, we will examine a simple procedure for estimating the underlying parameters of a simulated data set.

To generate the dataset, we will assume the simple model of tumour suppressor loss from above (see figure 1). To simulate the appearance of a tumour in a reference population of size *N, N* runs are performed with the Gillespie algorithm, with a known set of model parameters *θ*_*true*_ = {*α*_*j*_, *β*_*j*_, *κ*_*j*_, *µ*_*jk*_}, and the initial condition that every cell is initially on node 0. Cells on nodes 3 or 4 are neoplastic. The biological interpretation of the end node is whether or not the tumour presents with clonal LOH. In a more general model, the end node can represent what pattern of copy-number alterations (CNAs) is clonal in the resulting neoplasm [3, 18].

**Figure 1:**
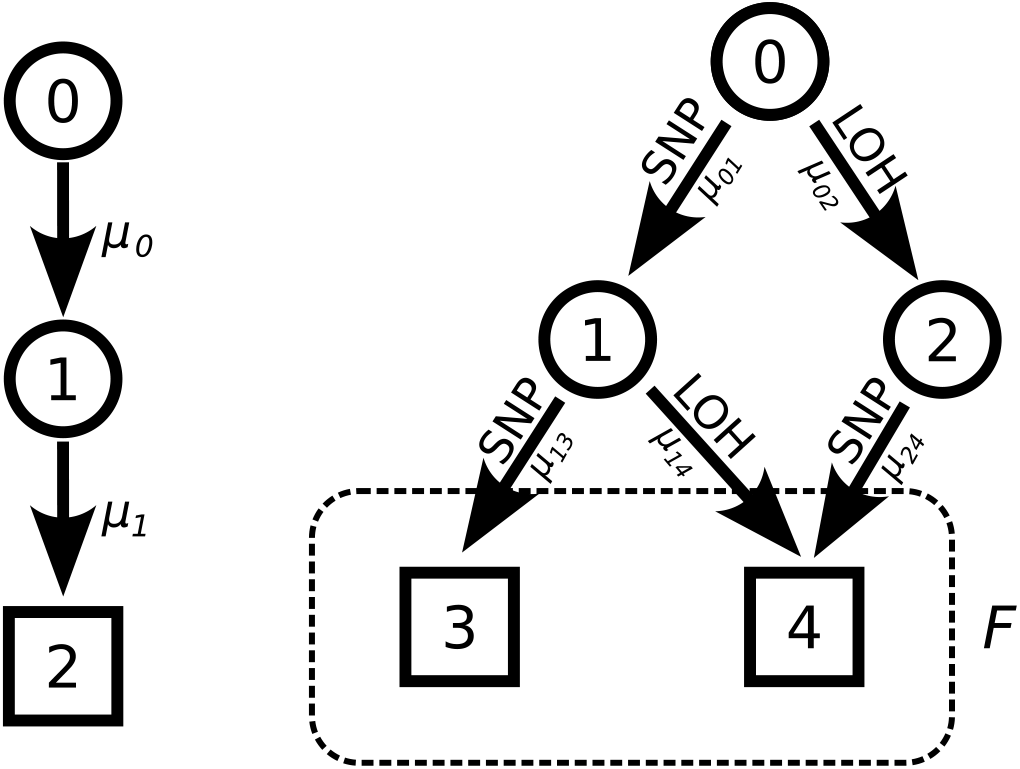
Left: an example of a two-step multi-stage clonal expansion model (MSCE) [2, 7]. Right: a generalised MSCE on a directed acyclic graph with five nodes, with the end nodes (squares) in the set *F* labeled. This shows how two different mechanisms of tumour suppressor loss can be resolved in a graphical model. The two mechanisms in question are single-nucleotide variants on different alleles, and copy number loss on the locus. The different end nodes in the set *F* correspond to different clonal copy number alterations in the resulting tumour: tumours of type 3 will have two mutated copies of the tumour suppressor, whereas those of type 4 will have one mutated copy and loss of heterozygosity on that locus.

If, in a given run, no cells arrive at either final node 3 or 4, this patient is considered not to have developed a primary tumour, and the age and type are not recorded. Thus, only a subset *m* of the *N* individuals in the simulated reference population will develop a tumour, and only *m < N* of the simulations will give a positive result.

The dataset therefore consists of a set of *m* observations, which consist of ages *t*_*i*_ and end nodes/tumour types *f*_*i*_, and the size *N* of the reference population. Once this dataset has been generated, the statistical problem is to find the best possible underlying parameters *θ* = {*α*_*j*_, *β*_*j*_, *κ*_*j*_, *µ*_*jk*_}. The most general way to achieve this is a maximum likelihood approach [12]. We will not attempt to infer the number of vertices on the underlying graph *G*, or the initial population *Z*_0_.

Once generated, the simulated dataset can be represented as a set of histograms, with one histogram for each tumour type *f* (see figure 2). In each histogram, the data i s binned according to age. The range of ages is split into uniform bins of width Δ*t*: the *i*^*th*^ bin has a start age *t*_*i*_, and a count *k*_*i*_. To compute the likelihood of this set of observations, a curve *S*_*f*_ (*t*) is computed given a set of parameters *θ*. The probability (or discrete hazard) *h*_*fi*_ that an individual will develop a primary tumour of type *f* in the *i*^*th*^ bin, which covers the interval [*t*_*i*_, *t*_*i*_ + Δ*t*], is

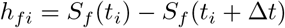

**Figure 2:**
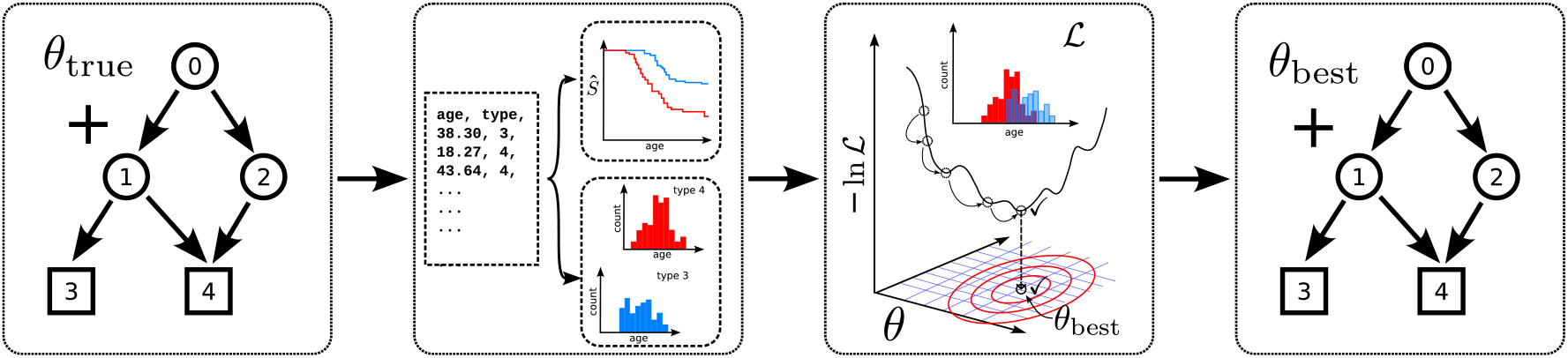
A figure describing the software framework that tests parameter inference by maximum likelihood estimation on a simulated longitudinal study. 1: A ground-truth model is proposed with known parameters *θ*_*true*_, and the Gillespie algorithm is used to generate a set of ages and types of diagnosed primary tumours, simulating a longitudinal clinical study. 2: the data set can be represented either with Kaplan-Meier estimators *Ŝ*(*t*) or a set of incidence histograms *I*_*k*_(*t*_*i*_). 3: the incidence in the simulated study is used to evaluate a likelihood function −ln ℒ(*θ*) with our new fast forward method. This likelihood is then minimised using a combination of brute force and gradient descent, although other approaches are possible. 4: the final output of the harness is a set of estimated parameters *θ*_*best*_, which may then be compared to the ground truth parameters *θ*_*true*_. See section 3.

If *n*_*i*_ ≤*N* individuals in a reference population of *N* individuals have survived to the *i*^*th*^ bin without being diagnosed with a tumour of type *f*, the number *k* that develop a tumour of type *f* in between the ages of *t* and *t* + Δ*t* will be binomially distributed. Following S. Johansen [34], the likelihood ℒ of observing *k*_*i*_ cases in the *i*^*th*^ age bin given *θ*, which includes ages between *t*_*i*_ and *t*_*i*_ + Δ*t*, is

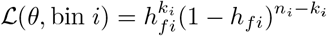

up to an irrelevant combinatorial prefactor. The combined likelihood for the simulated dataset is therefore

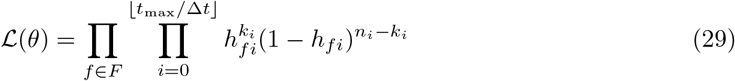

which corresponds to the likelihood for the Kaplan-Meier estimator *Ŝ* [34].

As a proof-of-concept, we can couple our one-pass fast forward method with a simple numerical optimisation method to maximise the above likelihood: to test if it can infer the parameters *θ*_*true*_ used to generate a synthetic dataset. The relevant *S*_*f*_ curves are computed numerically for each candidate set of parameter values *θ* using the numerical method from section 2.4. Briefly, after choosing a good start point through an initial round of brute force exploration, the parameters *θ* are varied using gradient descent, with the objective function defined to be −lnℒ from (29). Explicitly:

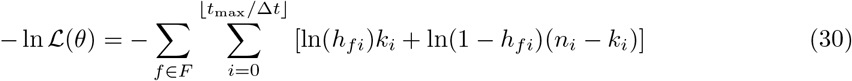

The best possible estimate found using this method (the “best guess”) will be denoted *θ*_*best*_.

## 3 Results

### 3.1 Scaling of errors and speed with step size Δ*t*

In the following, we will define the error to be the root mean squared error a long a given time interval *t* ∈ [0, *t*_max_].

The value of a fast numerical scheme for estimating *S*_*f*_ (*t*) immediately becomes apparent when comparing the run-time to random sampling, the only method known to be exact for birth-death processes on arbitrary graphs. To compare the methods fairly, we must determine how many replicates for Gillespie is equivalent to a chosen time step for the new, fast forward method. That is, the error in both methods should be on the same order of magnitude.

Random sampling estimates the underlying probability *P* of some event by performing *N* simulations, and counting the number of simulations *k* in which the event occurred. The estimate of *P* is then just *P* = *k/N*. As the samples are independent and identically distributed, the count *k* will be binomially distributed, and will have an associated variance

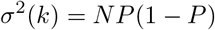

The associated standard deviation in the estimate *P* = *k/N* is therefore

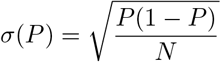

and thus we expect the error to scale as *ϵ*_*G*_ = 𝒪(*N*^−1*/*2^). This is indeed supported by simulations (figure 3).

**Figure 3:**
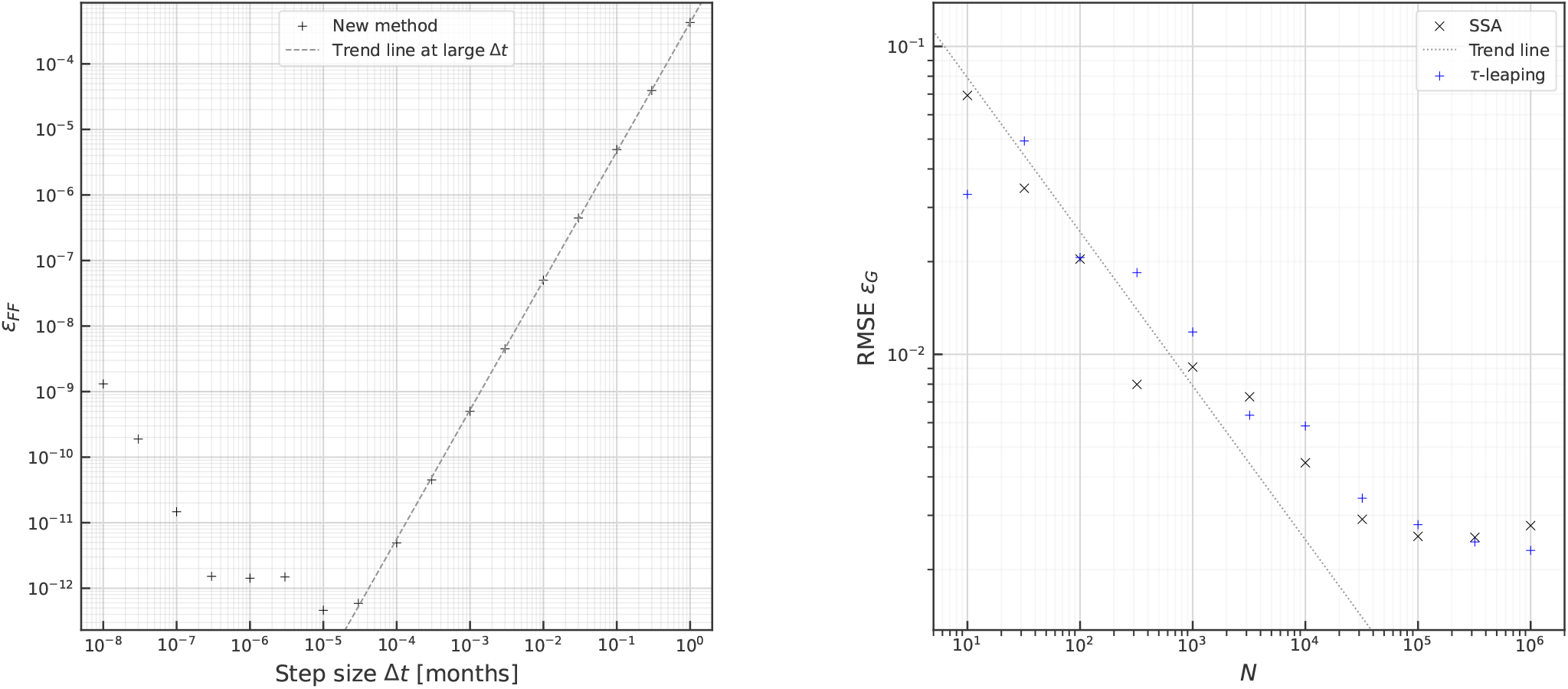
Plots showing error dependence of the Gillespie algorithm and our novel one-pass “fast forward” algorithm detailed in section 2.4. The model simulated was the 5-node model of tumour suppressor loss depicted in figure 1. Left: The scaling of the global numerical errors in the “fast forward” method described in 2.4 as a function of step size Δ*t*. The “floor” associated with round-off error is visible on the right, and on the left the upward 𝒪(Δ*t*^2^) scaling of truncation error is apparent as Δ*t* increases: the dashed line is the regression curve *ϵ*_*FF*_ ≈ 1.27 × 10^−3^ Δ*t*^1.97^. The global errors were estimated using Richardson extrapolation, comparing the computed value using a step size Δ*t* to one at 0.5Δ*t* [35]. Right: The scaling of global numerical errors for the Gillespie algorithm, defined as the mean squared difference between *Ŝ*_*k*_ for two simulations with different seed values. The dotted line is 0.5*N*^−1*/*2^, estimated in section 3.1.

The global error in a fast forward method that uses a numerical scheme of order-*p* will be *ϵ*_*F F*_ = 𝒪(Δ*t*^*p*^) [35, 36]. Because the local error in Heun’s method is 𝒪(Δ*t*^3^), the global error is 𝒪(Δ*t*^2^). That is, *p* = 2, and Heun’s method is second-order [36]. Thus, the global error *ϵ*_*F F*_ in the fast forward method’s estimate for *S*_*f*_ (*t*) should be

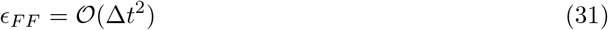

Numerically, the global error in a solver of ordinary differential equations can be estimated using Richardson extrapolation [35, 37, 38, 39]. This compares a solution for *P*(*t*) obtained using a given step size Δ*t* to the value obtained using a step size of Δ*t/*2 at the same time *t*, and forming a linear combination of the two values [38]. This clearly shows that the truncation error associated with the finite step size Δ*t* scales as 𝒪(Δ*t*^2^) as expected (figure 3).

For a fair comparison of the two techniques of computing *S*_*f*_ (*t*), we should choose Δ*t* so that the targeted global error size *ϵ* for the fast forward method is the same order of magnitude as the global error for a set of *N* Gillespie algorithm simulations. That is,

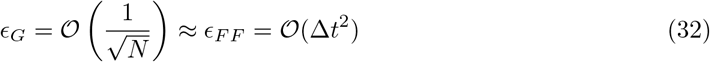

The run time *T*_*G*_ of the set of Gillespie algorithm simulations should be simply proportional to *N*, the number of runs performed, and so *T*_*G*_ = 𝒪(*N*) ∼ 𝒪(*ϵ*^−2^). On the other hand, the run time *T*_*F F*_ of one simulation with a fast forward method will scale as *T*_*F F*_ = 𝒪(1*/*Δ*t*) ∼ 𝒪(*ϵ*^−1*/*2^). So, for a given error tolerance *ϵ*, we can expect

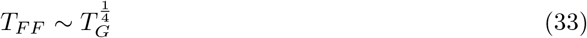

and for possible generalisations of the fast forward method that use order-*p* numerical integration schemes, it should be expected that

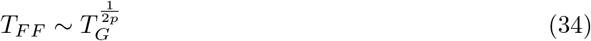

so that the run time of the fast forward method *T*_*F*_ will scale with a small power of the run time of the Gillespie algorithm *T*_*G*_. The one-pass fast forward method described in section 2.4 is order *p* = 2, so this method will outperform sampling with the Gillespie algorithm as *T*_*F F*_ ∼ *T*_*G*_^1*/*4^. For example, if we wanted to reduce the error *ϵ* in *P*(*t*) by a factor of 1*/*100, *T*_*F*_ would have to increase by a factor of 10, but *T*_*G*_ would become 10^4^ times more costly.

Finally, we can repeat the above asymptotic analysis for Quinn’s proposal, which used Euler’s method as an integrator. Euler’s method has a global error *ϵ*_*Q*_ = 𝒪(Δ*t*), and Quinn’s scheme needed two passes, so the runtime *T*_*Q*_ is of order 𝒪(Δ*t*^−2^). Therefore, *T*_*Q*_ ∼ 𝒪(*ϵ*^−2^), which has the same scaling with target error *ϵ* as random sampling with Gillespie’s algorithm. In this sense, Quinn’s method is asymptotically just as slow as random sampling, which may explain why this approach was not seriously pursued.

### 3.2 Profiling and comparison of methods

For the Gillespie simulations, the survival curves *S*_*k*_ were estimated using Kaplan-Meier estimators. To estimate the error in the Gillespie approximation, two runs with the same sample size *N* were conducted using different seed values for the random number generator. The squared difference between the computed values for the resulting survival curves, *Ŝ*_1_ and *Ŝ*_2_, was rescaled by 1*/*2 and used as an estimate for the local mean squared error. The mean value of this local mean squared error was then used as a metric for the global error.

Both the “fast forward” simulations and stochastic simulations were carried out on the University of Manchester’s high-performance computing cluster CSF3 *“Danzek”*.

Figure 3 shows that the global errors in the two methods being compared scale as predicted in section 3.1. Furthermore, figure 5 shows that the scalings of runtime *T* with error tolerance *ϵ* of the two methods is also as predicted. It follows that the fast forward method will vastly outperform random sampling as the error threshold of interest *ϵ* is decreased, and this is indeed what is observed in figure 5.

**Figure 4:**
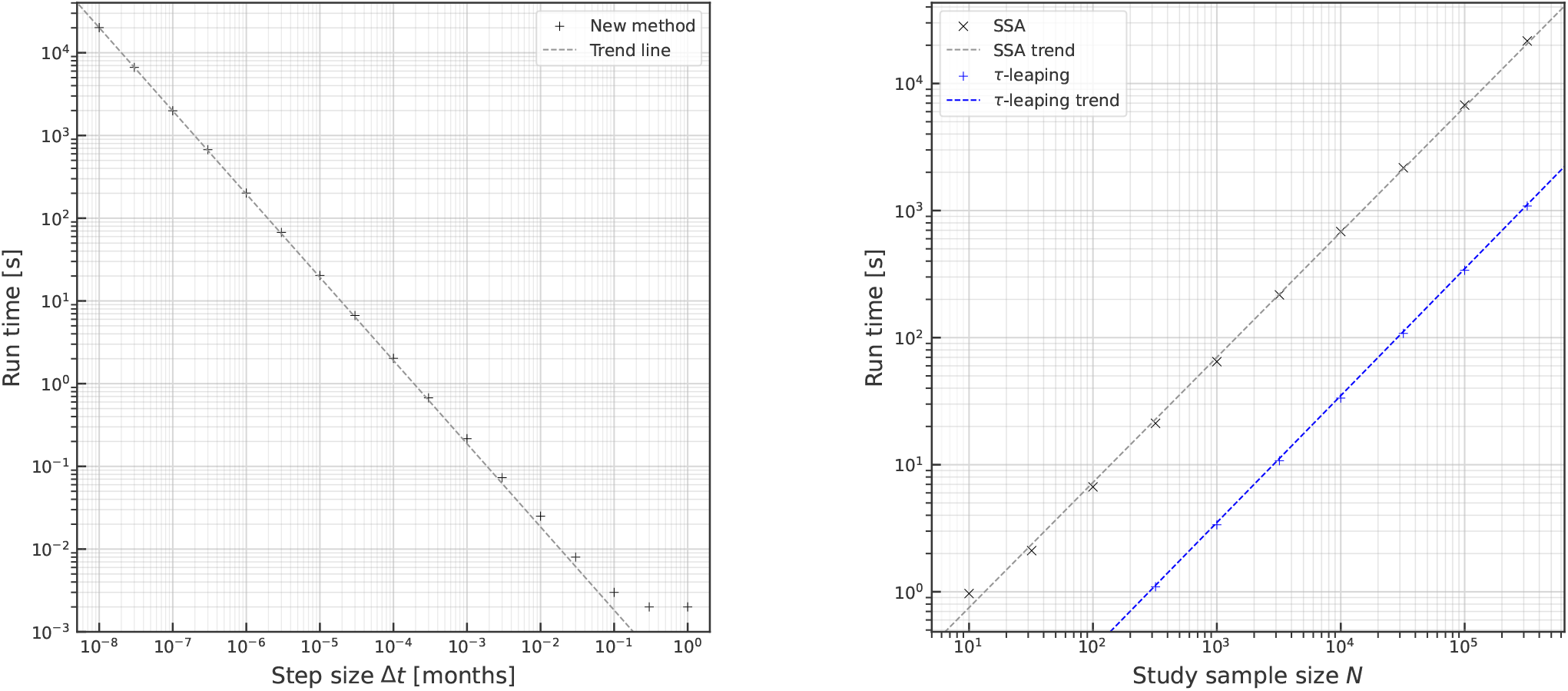
Plots of the real run time of the fast forward method as a function of Δ*t* (left), and the run time of the stochastic method (right). The regression lines are: left, black: *T* ≈ 1.80 × 10^−4^(Δ*t*)^−1.00^, consistent with our argument that *T* ∼ Δ*t*^−1^; right: trend lines for the Gillespie algorithm (black) and *τ*-leaping (blue). The Gillespie algorithm has *T* ≈ 7.77 × 10^−2^*N* ^0.985^, and *τ*-leaping has *T* ≈ 3.54 × 10^−3^*N* ^0.998^. This is also consistent with our argument that *T* ∝ *N*.

**Figure 5:**
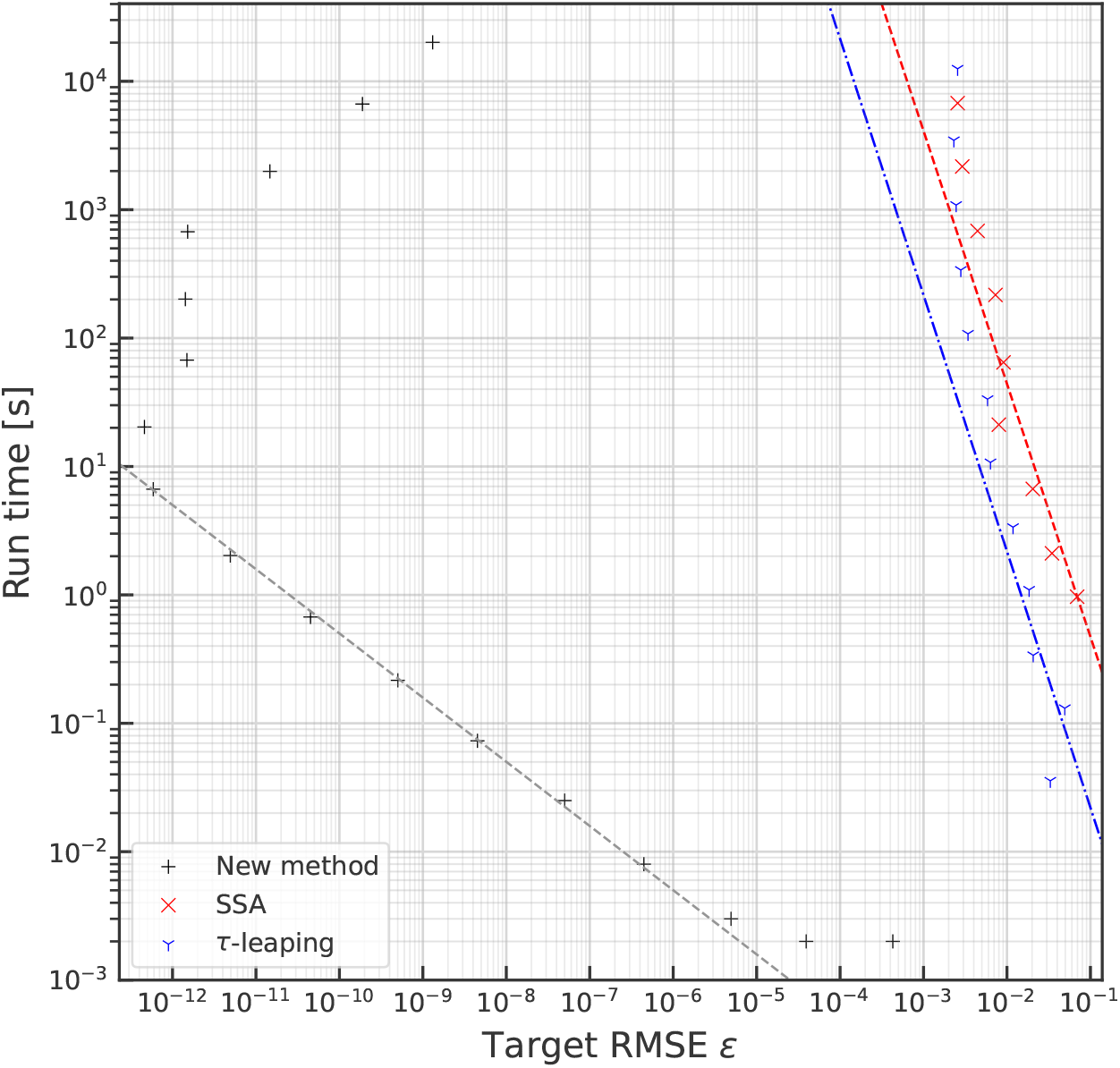
A plot showing how run time depends on desired root-mean squared error for both stochastic sampling with the Gillespie algorithm (red × s), *τ*-leaping (blue *Y* s), and our new fast forward method (black +s). The *T* ∼ *ϵ*^−1*/*2^ dependence of the fast forward method is apparent (black line), and a significant improvement on the *T* ∼ *ϵ*^−2^ scaling of random sampling with either Gillespie’s algorithm (red line) or *τ*-leaping (blue line). Even for *τ*-leaping, the fast forward method is 10^7^ times faster for an error tolerance of *ϵ* = 10^−4^, a dramatic improvement of seven orders of magnitude.

Comparing Kaplan-Meier estimators from Gillespie algorithm simulations to the *S*_*k*_-curves from the fast forward method (figure 6) shows excellent agreement across most of the range of the dataset, but reveals a surprising feature: the Kaplan-Meier and *S*-curves visibly diverge at late ages. Close inspection reveals that this is an artifact resulting from extrapolating the KM estimator beyond the range of the underlying data. As a non-parametric estimator, it cannot capture information about underlying trends that continue after the last item in the dataset, as these are fundamentally parametric in nature: in this case, the exponential fall-off in old age. As such, it is only valid to compare the errors in the two methods where data was available in the stochastic simulations. “Zooming in” on the problematic region confirms this: the numerical *S*-curve continues to decrease smoothly with age, whereas Kaplan-Meier estimators are flat beyond the oldest entry in the dataset. Increasing the number of replicates of the stochastic simulations improves agreement further, as visible in the detail on the right of figure 6.

**Figure 6:**
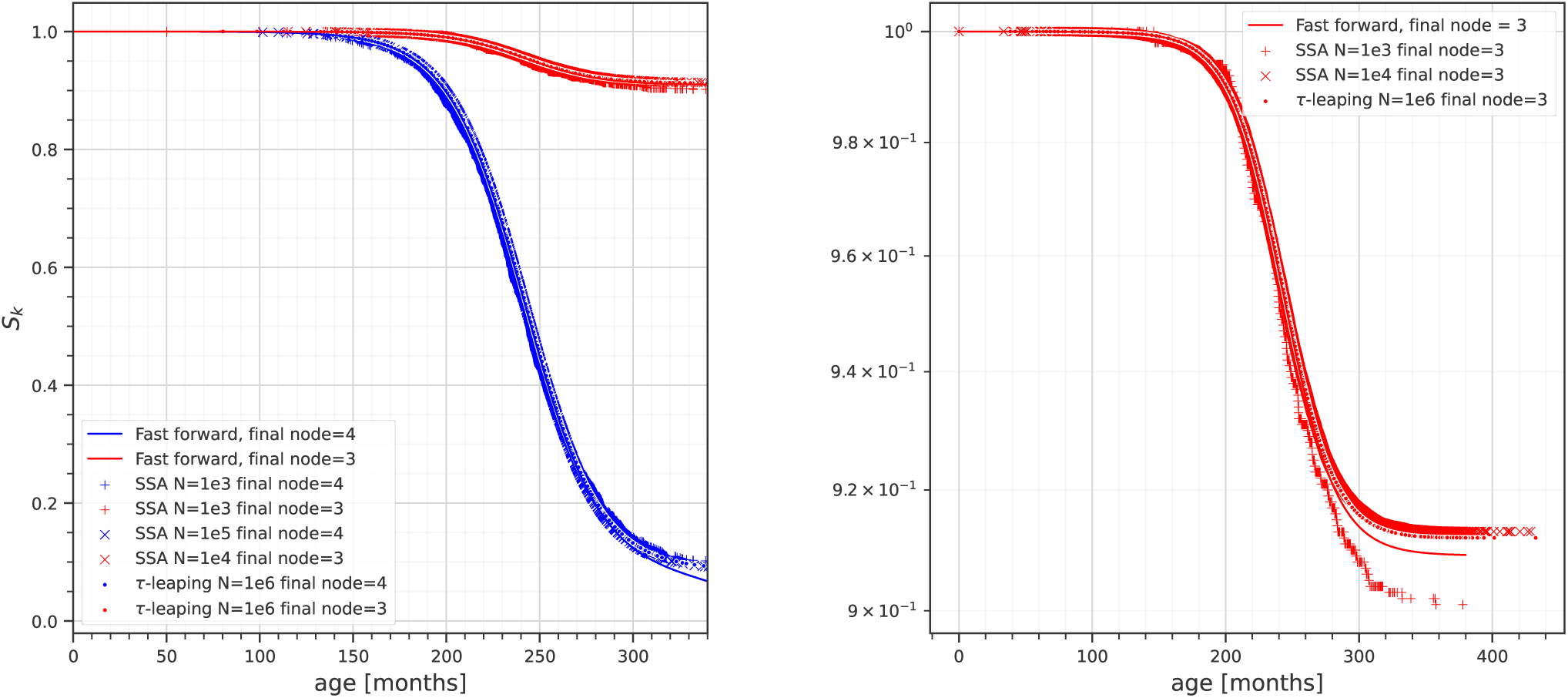
Left: a comparison of *Ŝ* curves for the tumour suppressor loss study generated by the Gillespie algorithm (+ signs: computed with *N* = 10^4^ replicates: · signs: computed with *N* = 10^6^ replicates), and the corresponding set of *S*_*k*_ curves generated by our fast forward algorithm from section 2.4 (dashed). Right, detail: an artifact causing deviations between the *Ŝ* and *S*_*k*_ curves at late ages decreases as the sample size used for the stochastic simulations increases.

### 3.3 Application to statistical inference

The test framework described in figure 2 was used to generate simulated clinical studies and then guess the underlying parameters for a range of different study sample sizes. The sample sizes considered varied from 10 to 10, 000 and were evenly spaced on a logarithmic scale. To gauge the robustness of the estimates, the parameter estimates for each simulated study were cross-validated using jack-knife resampling [40].

Our new “fast forward” method for evaluating *S*_*k*_ is efficient enough that the likelihood function ℒ from 29 becomes relatively cheap to evaluate, which allows us to gain highly detailed information about its structure in hypothesis space. As well as the best guess from maximising the likelihood, it is possible to compute a Fisher information matrix (from the Hessian of − logℒ) near that best guess, which was used to generate analytical conference intervals using Wilks’ theorem [41]. However, it should be remembered that the maximum and Fisher information matrix only adequately characterise the distribution if the matrix Is nonsingular and the distribution is not fat-tailed – in other words, if ℒ looks sufficiently like a normal distribution.

The main result of this part of the work Is that while maximum likelihood estimation does estimate values for the underlying parameters that are on the correct order of magnitude with minimal prior information, the deviation between the maximum likelihood estimates and the ground truth is stubbornly high, and converges very slowly as the sample size increases (see figure 7). This was a surprise. The underlying reason Is that the likelihood function ℒ is *non*-normal in a number of important ways, and is not well-behaved. We suspect that there Is an approximate symmetry in ℒ that relates *r*_*LOH*_ and *s*_2_. making this pair unidentifiable [42, 43].

**Figure 7:**
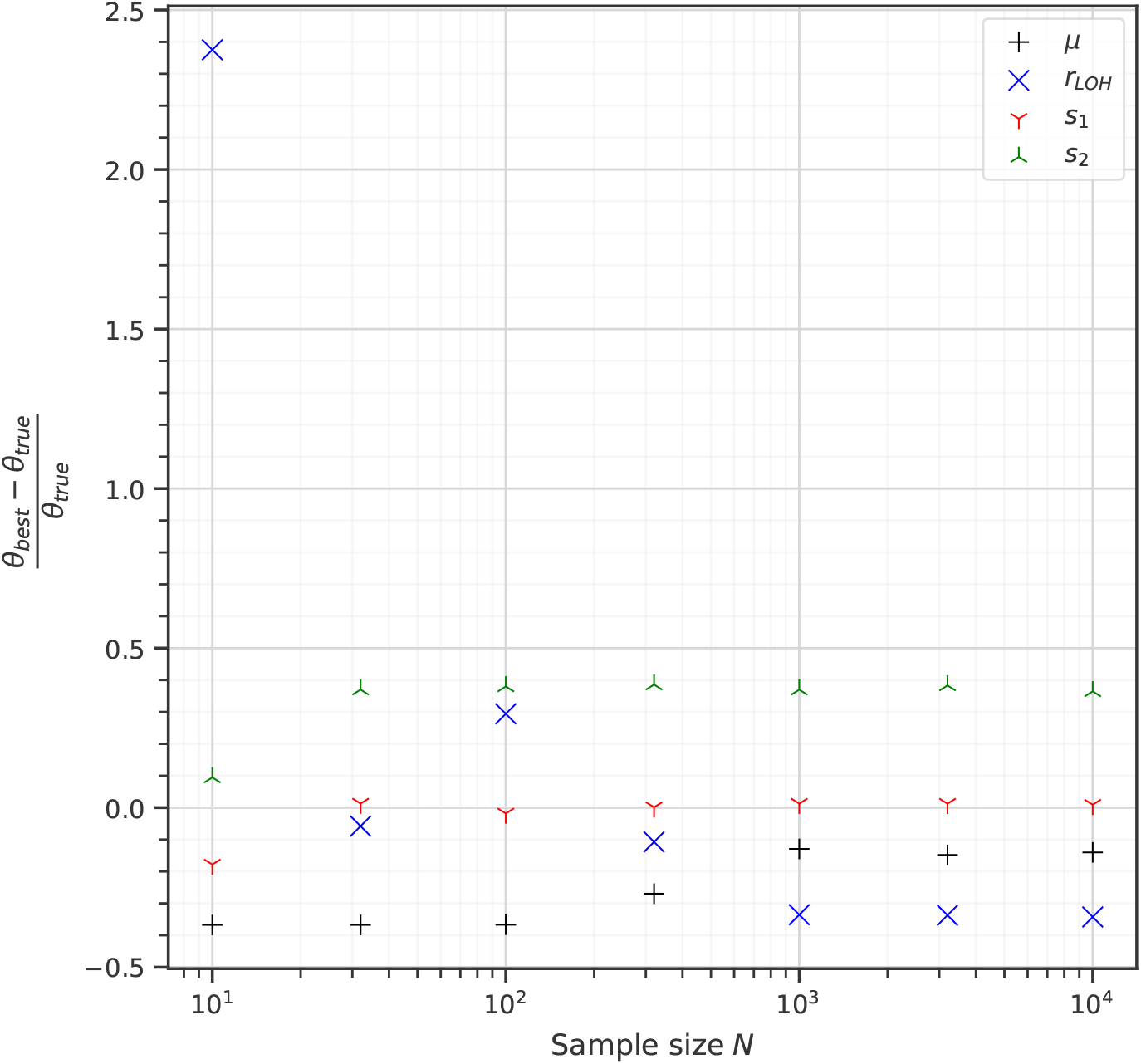
Figure showing relative errors in the output of the test harness by parameter: *µ*, black +s; *r*_*LOH*_, blue ×s; *s*_1_, red *Y* s; and *s*_2_, green ⊥s. While estimates of *µ* and *s*_1_ converge to the ground truth as the study sample size *N* increases, *r*_*LOH*_ and *s*_2_ remain trapped a local minimum, both about ±40% away from their real ground truth values in either direction. This was a surprise, and prompted a deeper investigation of the structure of the likelihood function − lnℒ.

But first, to avoid the known pitfalls of parameter inference in MSCE models, it must be noted that the initial precursor cell population iV_0_ is not identifiable separately of when *N0* Is very large, a problem extensively studied in prior work [10, 1, 43]. Put briefly, this follows from a symmetry in the model dynamics and the stoichiometry of cell division: when death of wild-type cells is zero, and the population growth rate also zero, the initial population will be constant, and only the product *μN*_0_ will be observable [10. 1). To address this, the population *N*_0_ Is fixed to a known value rather than inferred, and only the mutation rates *μ,r*_*LOH*_ and fitnesses *s*_1_,*s*_2_ were varied during likelihood maximisation.

Even after known was taken into account, problems with likelihood maximisation remained. The condition numbers computed from the Fisher information matrix were large (see figure 11), indicating that the discovered best guesses lay in a region that was close to being singular, with a small eigenvalue. The Fisher information matrix was therefore ill-conditioned, which may be due to a near-symmetry in the underlying model. No exact symmetries were apparent on analytical inspection [42].

**Figure 8:**
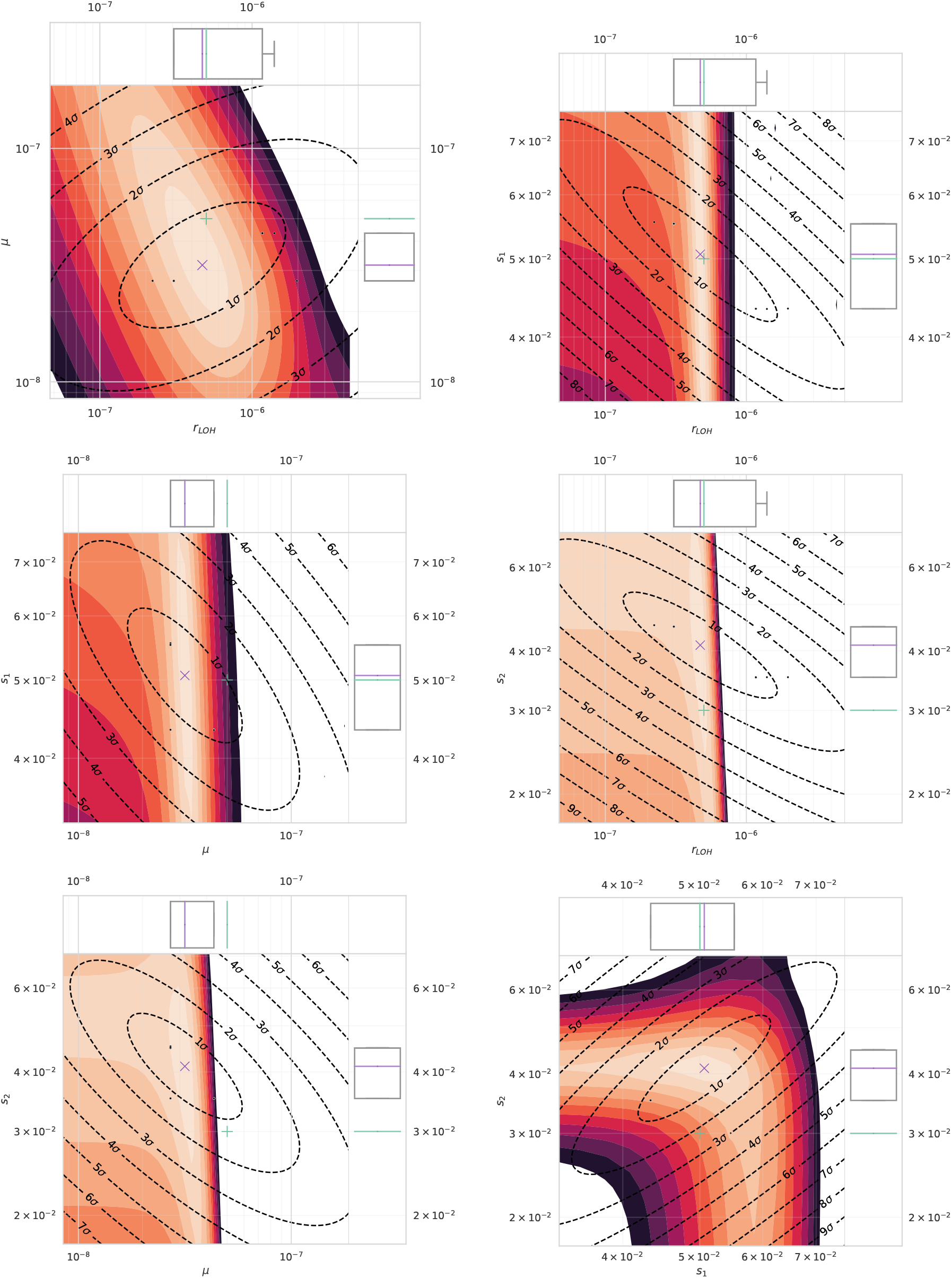
Results from the test harness, showing the parameter values for the ground truth (red +), best guess (blue x); a cross section of the likelihood function as a heat map (light orange to black), with likelihood levels that correspond to usual confidence intervals; confidence intervals derived from the Fisher information matrix (black-to-orange ellipses), and a point cloud representing resampled best guesses (grey circles). Top left: *µ* and *r*_*LOH*_ ; top right: *r*_*LOH*_ and *s*_1_; centre left: *µ* and *s*_1_; centre right: *r*_*LOH*_ and *s*_2_; bottom left: *µ* and *s*_2_; bottom right: *s*_1_ and *s*_2_. Sample size *N* = 32.

**Figure 9:**
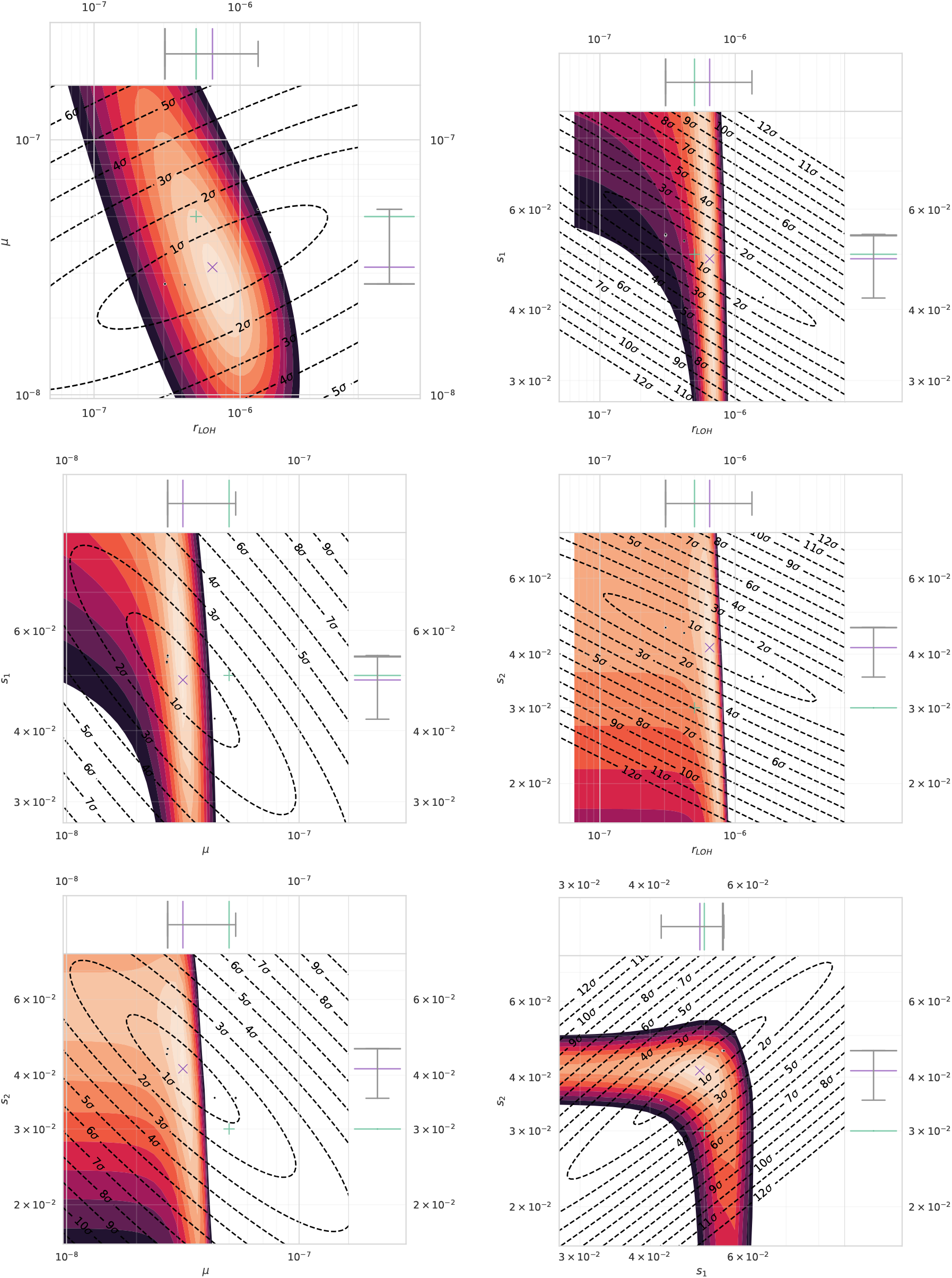
Results from the test harness, showing the parameter values for the ground truth (red +), best guess (blue x); a cross section of the likelihood function as a heat map (light orange to black), with likelihood levels that correspond to usual confidence intervals; confidence intervals derived from the Fisher information matrix (black-to-orange ellipses), and a point cloud representing resampled best guesses (grey circles). Top left: *µ* and *r*_*LOH*_ ; top right: *r*_*LOH*_ and *s*_1_; centre left: *µ* and *s*_1_; centre right: *r*_*LOH*_ and *s*_2_; bottom left: *µ* and *s*_2_; bottom right: *s*_1_ and *s*_2_. Sample size *N* = 100.

**Figure 10:**
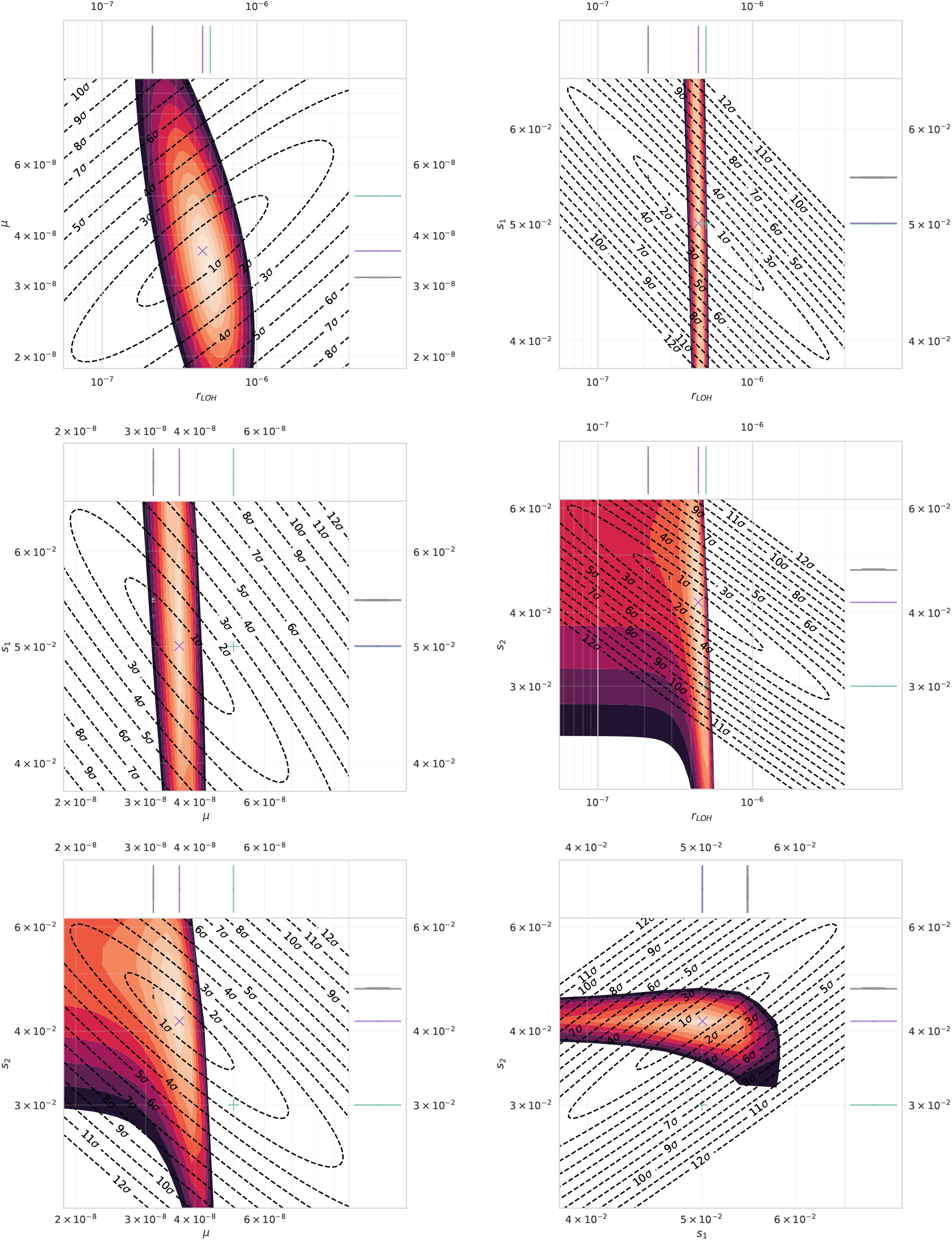
Results from the test harness, showing the parameter values for the ground truth (red +), best guess (blue x); a cross section of the likelihood function as a heat map (light orange to black), with likelihood levels that correspond to usual confidence intervals; confidence intervals derived from the Fisher information matrix (black-to-orange ellipses), and a point cloud representing resampled best guesses (grey circles). Top left: *µ* and *r*_*LOH*_ ; top right: *r*_*LOH*_ and *s*_1_; centre left: *µ* and *s*_1_; centre right: *r*_*LOH*_ and *s*_2_; bottom left: *µ* and *s*_2_; bottom right: *s*_1_ and *s*_2_. Sample size *N* = 320.

**Figure 11:**
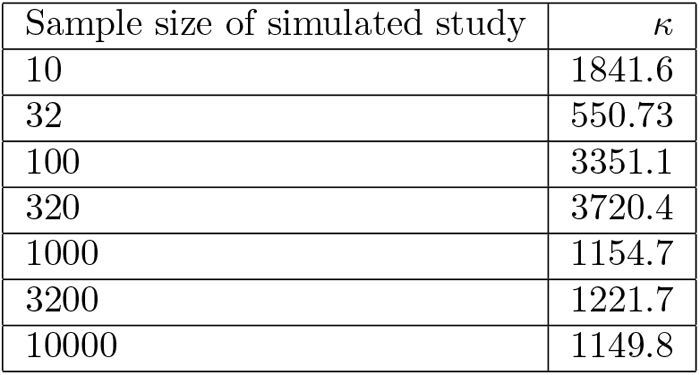
A table of condition numbers for each of the simulated clinical studies in the neighbourhood of the best guess. The condition number is defined as the ratio of the absolute values of the largest and smallest eigenvalues of the Fisher information matrix, *κ* := max_*i*_(|*λ*_*i*_|)/min_*i*_(|*λ*_*i*_|). Condition numbers this high indicate issues with identifiability, we believe due to an approximate symmetry.

Instead of simply maximising the likelihood and studying local properties of ℒ. the function was sampled in detail in a neighbourhood around the best guess. This gave a high-quality set of tests that revealed the structure of ℒ in detail. For each of the simulated clinical studies and estimates, six plots were made that showed the estimated and ground truth parameters with cross sections through ℒ. These plots revealed much fatter tails in ℒ than a normal distribution would have, as well as a long and narrow ridge in the likelihood function (explaining the high condition numbers and small eigenvalues), and strong correlations between the different estimates. These plots are shown in figures 8 - 10, including both confidence intervals derived from the Fisher information matrix and equivalent intervals in the likelihood function ℒ. These plots each show six cross-sections through the best guess *θ*_*best*_. Each cross-section shows how the likelihood varies with two of the four parameters.

Due to the issues with identifiability just mentioned, the plots for sample sizes of *N* = 10 and *N >* 1000 looked no more informative than the plots for intermediate sample sizes. For this reason the tests of the inference harness with sample sizes *N* = 32, 100, 320 are included in this manuscript as figures 8 - 10. The other plots are presented as figures 12 - 15 in the supplemental material.

**Figure 12:**
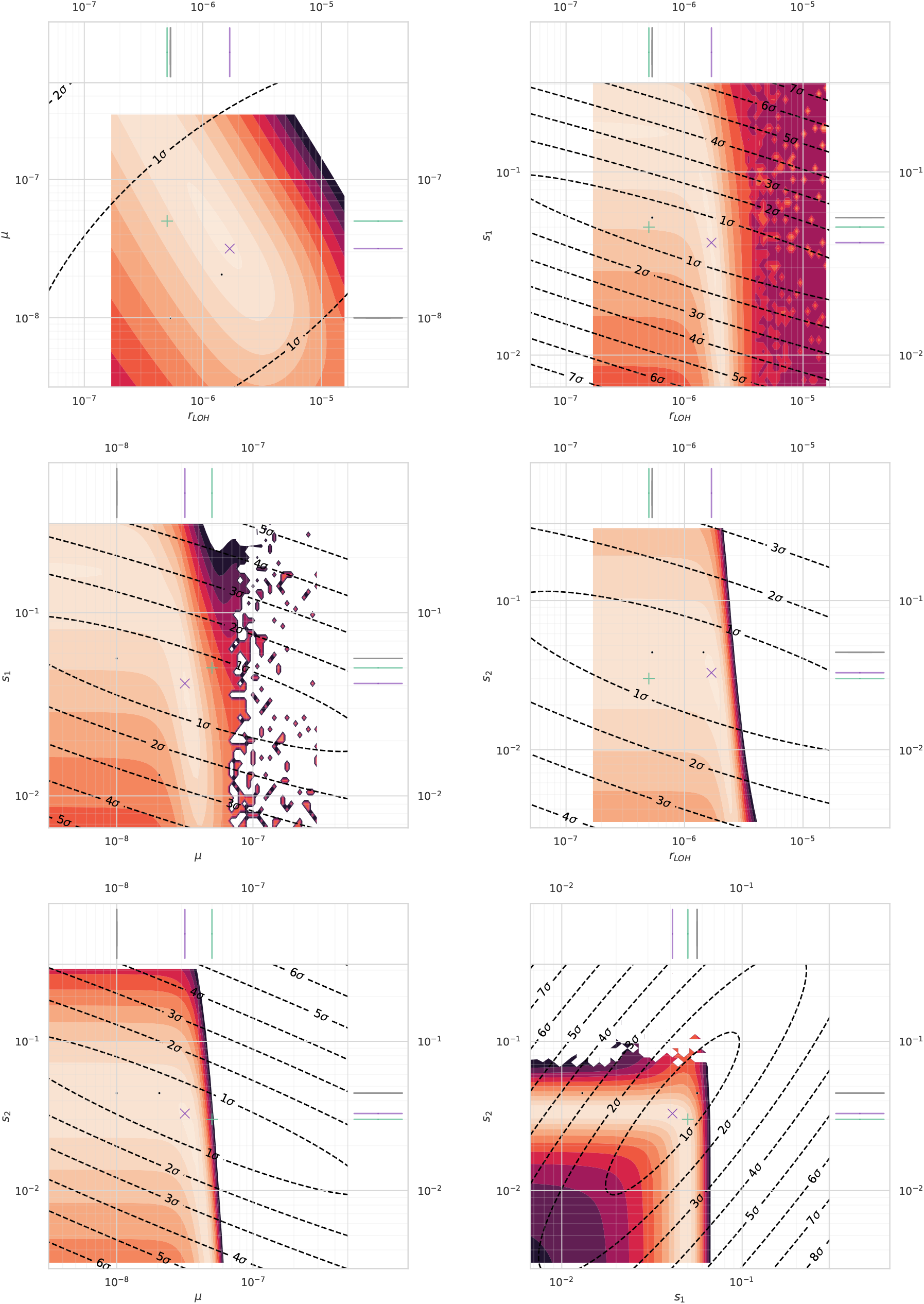
Results from the test harness, showing the parameter values for the ground truth (red +), best guess (blue x); a cross section of the likelihood function as a heat map (light orange to black), with likelihood levels that correspond to usual confidence intervals; confidence intervals derived from the Fisher information matrix (black-to-orange ellipses), and a point cloud representing resampled best guesses (grey circles). Top left: *µ* and *r*_*LOH*_ ; top right: *r*_*LOH*_ and *s*_1_; centre left: *µ* and *s*_1_; centre right: *r*_*LOH*_ and *s*_2_; bottom left: *µ* and *s*_2_; bottom right: *s*_1_ and *s*_2_. Sample size *N* = 10.

**Figure 13:**
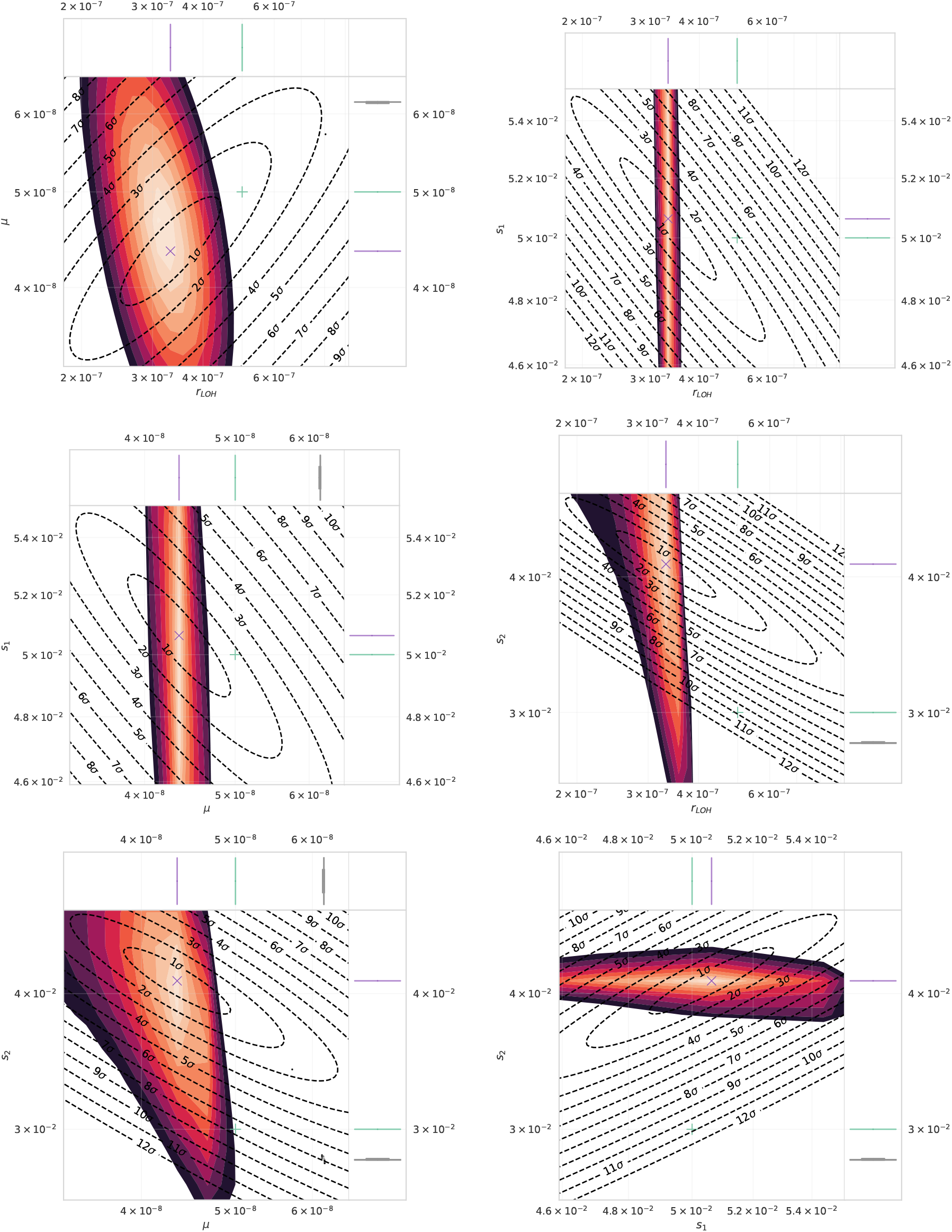
Results from the test harness, showing the parameter values for the ground truth (red +), best guess (blue x); a cross section of the likelihood function as a heat map (light orange to black), with likelihood levels that correspond to usual confidence intervals; confidence intervals derived from the Fisher information matrix (black-to-orange ellipses), and a point cloud representing resampled best guesses (grey circles). Top left: *µ* and *r*_*LOH*_ ; top right: *r*_*LOH*_ and *s*_1_; centre left: *µ* and *s*_1_; centre right: *r*_*LOH*_ and *s*_2_; bottom left: *µ* and *s*_2_; bottom right: *s*_1_ and *s*_2_. Sample size *N* = 1000.

**Figure 14:**
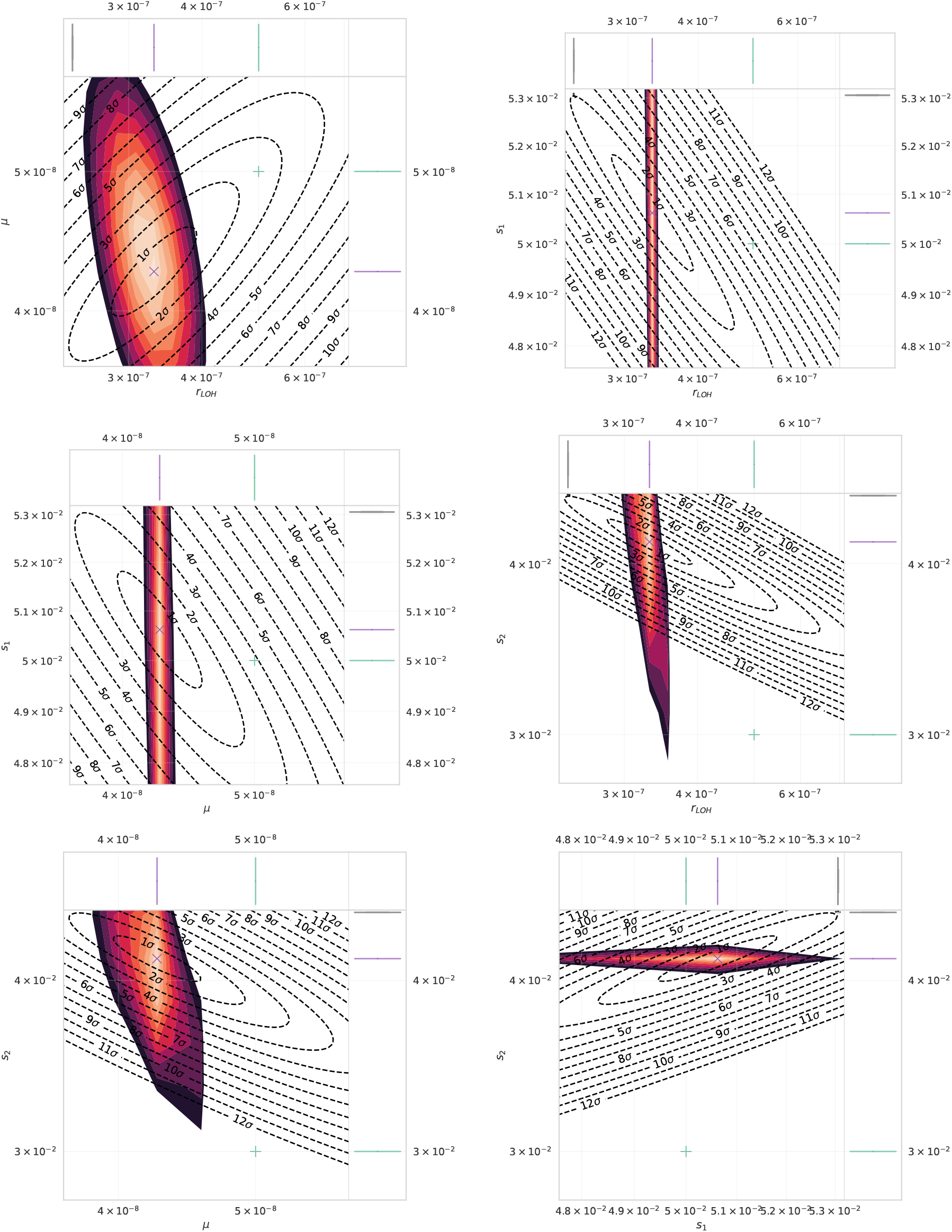
Results from the test harness, showing the parameter values for the ground truth (red +), best guess (blue x); a cross section of the likelihood function as a heat map (light orange to black), with likelihood levels that correspond to usual confidence intervals; confidence intervals derived from the Fisher information matrix (black-to-orange ellipses), and a point cloud representing resampled best guesses (grey circles). Top left: *µ* and *r*_*LOH*_ ; top right: *r*_*LOH*_ and *s*_1_; centre left: *µ* and *s*_1_; centre right: *r*_*LOH*_ and *s*_2_; bottom left: *µ* and *s*_2_; bottom right: *s*_1_ and *s*_2_. Sample size *N* = 3200.

**Figure 15:**
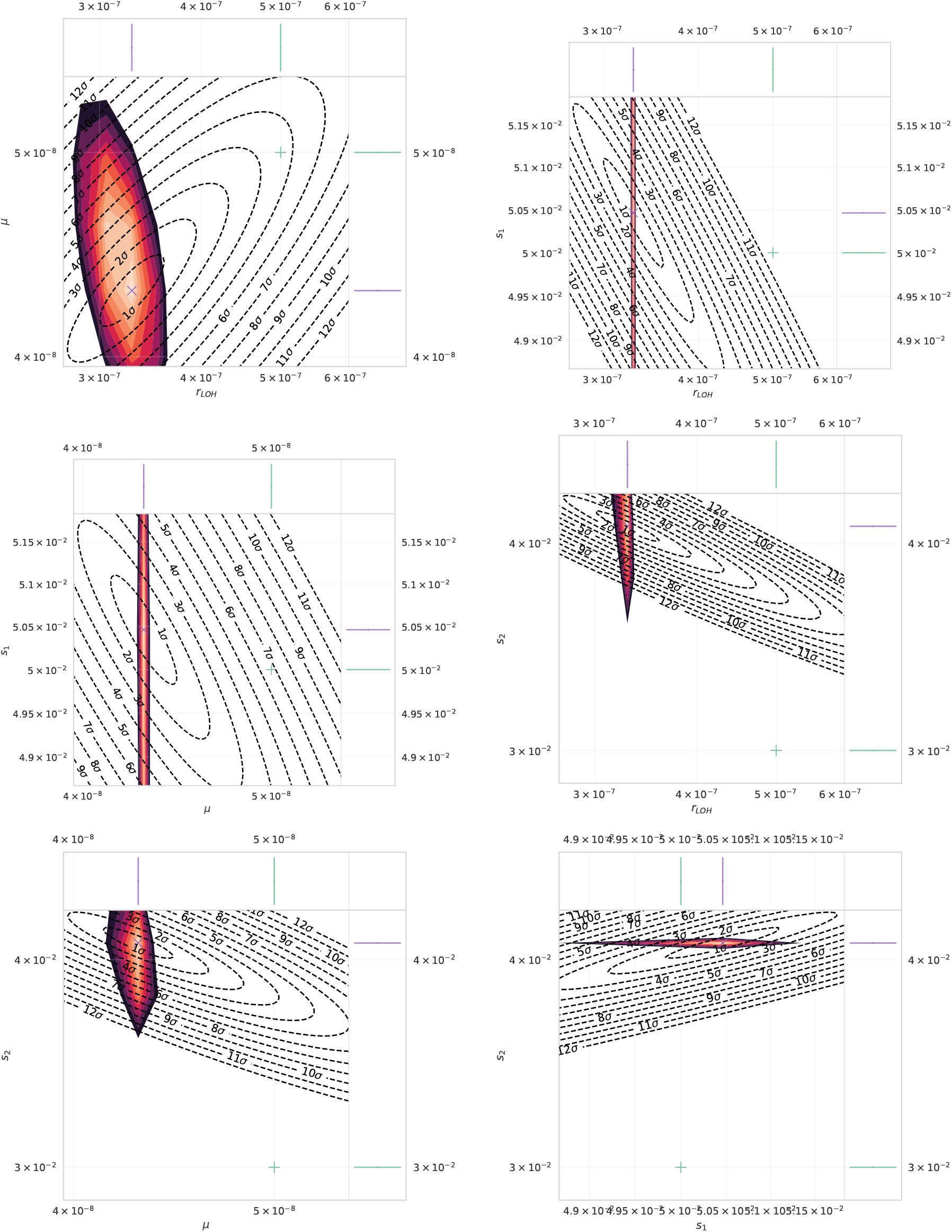
Results from the test harness, showing the parameter values for the ground truth (red +), best guess (blue x); a cross section of the likelihood function as a heat map (light orange to black), with likelihood levels that correspond to usual confidence intervals; confidence intervals derived from the Fisher information matrix (black-to-orange ellipses), and a point cloud representing resampled best guesses (grey circles). Top left: *µ* and *r*_*LOH*_ ; top right: *r*_*LOH*_ and *s*_1_; centre left: *µ* and *s*_1_; centre right: *r*_*LOH*_ and *s*_2_; bottom left: *µ* and *s*_2_; bottom right: *s*_1_ and *s*_2_. Sample size *N* = 10000.

There are apparent saddle points in the cross-sectional plots, which can indicate multiple local extrema. To study this, ℒ was sampled in a larger neighbourhood of the best guess, and volumetric ray casting was used to produce detailed 3-dimensional video renderings (included in the supplemental material). These renderings map *µ, r*_*LOH*_, and *s*_1_ to the 3 spatial dimensions, mapping values of *s*_2_ to colour, and used normalised probabilities ℒ to indicate brightness [44]. The results revealed a complex, non-monotonic function that, in addition to the long narrow ridge that contained the best guess, had curious branching structures. As visible in the video renderings, the central, long, narrow ridge appears to double back on itself, with at least one additional maximum at high values of *s*_1_ and *s*_2_.

Despite the difficulties presented by these complex structures in the likelihood function, it can be seen from figures 8-10 (and supplemental figures 12-15) that the best guess was close to the ground truth in most cases, and agreed adequately even for relatively small sample sizes of *N* = 10 and *N* = 32. This suggests that the method holds considerable promise for inferring mutation rates and other genomic parameters in relatively rare diseases, where sample sizes are limited, as it can provide new, strong constraints from combined genomic and demographic data making minimal assumptions about initial values. However, identifiability remains a problem. The long, narrow ridge and other structures in ℒ mean that there are large asymmetries in the confidence intervals, and not all of the parameters are identifiable, which future research must try to address or mitigate.

## 4 Discussion

The fast algorithm presented in section 2.4 allows survival curves and first-passage times for clonal selection models to be evaluated rapidly and deterministically, seven orders of magnitude faster than any practical method based on random sampling. This in turn enables mutation rates and fitness advantages during carcinogenesis to be estimated from simulated data using maximum likelihood methods, without the use of approximate Bayesian computation or related techniques. This computation of likelihood functions was not previously feasible. However, although the likelihood function for this model can be computed, much additional work remains to improve the identifiability of the model. We have shown using simulated clinical studies that the resulting risk models should be able to combine epidemiological and genomic data, correlating age with copy number alterations, or other genomic rearrangements. In the future, these models may be able to capture individualised risk factors: perhaps showing how rare variants or genomic signatures interact with patient age.

This class of model should permit better estimates of parameters of biological mechanisms, as well as improved forecasts of cancer incidence with age and risk factors, with clear applications in public health modelling. [3, 4, 16]. While the method described here was motivated by models of cancer incidence, the numerical scheme should also be applicable to any chemical reaction network that is first-order and has constant rate coefficients [27]. Although the “fast forward” method described in section 2.4 has key limitations – namely, for all reactions to be first order, and for the coefficients to be constant – it is clearly applicable to reactions with more than one product, which gives it a broader scope than known exact solutions [19, 21, 22, 23].

It is also interesting to note that the various “classical” methods based on Kolmogorov backward equations typically have several generating functions Φ_*k*_ arranged in a hierarchy [1, 15, 14]. In contrast, both our “fast forward” method from section 2.4 and Quinn’s earlier fast forward method rely on only one generating function Ψ with multiple arguments. The analogous variables to Φ_*k*_ in our approach are the characteristic curves, *γ*_*k*_ [8].

Our new method is limited to models with constant coefficients. Continued work must discover an efficient numerical integration scheme for evolution on graphs that is suitable for time-varying parameters *θ*(*t*) = {*α*_*j*_(*t*), *β*_*j*_(*t*), *µ*_*jk*_(*t*), *κ*_*j*_(*t*)}. This will be necessary to study carcinogenic processes associated with mutagen exposure, chronic inflammatory conditions, or other environmental factors. These are relevant to many important public health challenges, such as Barrett’s oesophagus, obesity, and smoking [2, 12]. Quinn’s algorithm of 1988 is one candidate, but our analysis in section 3.1 and contemporary reviews suggest that it is asymptotically not very efficient [13, 11, 10]. On the other hand, work has shown that there may be a family of unexplored relatives to this algorithm with considerable optimisations, both regarding the rate of convergence of the numerical integrator and the amount of necessary caching. A two-pass method like Quinn’s that used an order-*p* integrator would have a theoretical runtime complexity of 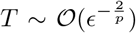, so further gains over random sampling should be possible if a higher-order Runge-Kutta scheme were used.

It is also interesting to note some similarities between the graphical models described here and neural networks. Like neural networks, the models have a graph structure, and the edges *E* and nodes *V* carry “weights” given by the model parameters *θ* and subpopulations 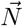. The survival curve *S*_*f*_ (*t*) is analogous to a neural network’s activation function. These similarities to neural networks raise the natural question of whether this class of models is differentiable, and if so whether backpropagation can be applied to maximise likelihoods with gradient-based methods.

Unlike neural networks, *S*(*t*) is not formed by simple function compositions; the populations 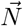 on each node are random variables; and the model parameters *θ* = {*µ*_*jk*_, *α*_*j*_, *β*_*j*_} have direct biological interpretations. The “weights” of this class of model are thus both radically different from typical neural networks, and have clear mechanistic interpretations. These additional subtleties make this class of model “explainable” or “interpretable”, in contrast to approaches that treat the model and its parameters as a black box [45, 46].

In short, we have established that Kolmogorov forward equations provide a viable approach for estimating hazard functions and likelihoods. Our work also suggests that a larger family of “fast forward” methods may exist for studying stochastic processes on graphs, which may prove a flexible alternative to existing methods based on Kolmogorov backward equations. Similarly, the algorithm used for maximum likelihood estimation in section 3.3 could be replaced with many other optimisation algorithms. The choice of which numerical integration scheme to pair with which algorithm to best infer the parameters of cancer evolution from genomic epidemiology is a new, open question.

## 5 Acknowledgements

The authors would like to acknowledge the assistance given by Research IT and the use of the Computational Shared Facility at The University of Manchester.

We would especially like to thank Ramishka Dona Liyanage for her work on the simulation codebase and the Python API; we would also like to thank Mark Kot and Greg Tumolo for helpful discussions about data visualisation and computer graphics; and also John C Baez, Amit Hazi, Clare Gratrex and Ruibo Zhang for engaging discussions regarding both the method and connections to related literature.

## A Appendix 1 definition of survival curve

In the case that the set of final nodes *F* has only one member, the relationship to the survival curve *S*_*f*_ (*t*) is straightforward, and

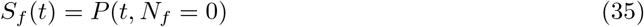

But in general, the set of final nodes *F* on the graph *G* may have more than one member. The model admits cases in which more than one subpopulation *N*_*f*_, *N*_*f*_*′* is nonzero simultaneously. Biologically, this corresponds to a scenario in which a tumour of type *f′* arises after a primary tumour of type *f* is already present (or vice versa). This is not unrealistic, as multifocality is observed in some types of brain tumour [47]. However, it complicates the definition of *S*_*f*_(*t*), as the simple probability *P*(*t, N*_*f*_ = 0) will include cases that are not primary tumours.

Recall that *S*_*f*_ (*t*) should be the probability that a patient survives to age *t* without a diagnosis of a primary tumour of type *f*. Without loss of generality, we can assume that the probability that tumours of two distinct types *f* ≠ *f′* cannot arise simultaneously. The relevant probability, so that we count only primary tumours of type *f*, is the probability that the subpopulation *N*_*f*_ = 0, conditioned on all other final subpopulations being zero:

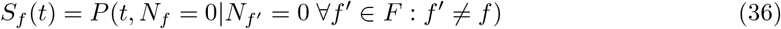

This counts only cases of type *f* where there is no existing tumour of type *f′* ≠ *f*. These cases are precisely the primary tumours of type *f*.

By the definition of conditional probability *P*(*A*|*B*) := *P*(*A* ∩ *B*)/*P* (*B*), equation (36) implies

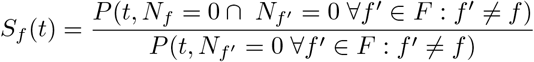

Which is equivalent to

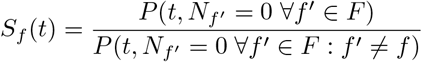

That is, compute *P*(*t, N*_*f*_*′* = 0, …) for all final nodes in *F*; then compute *P*(*t, N*_*f*_*′* = 0, …) for all final nodes *f′ except for f*; and *S*_*f*_ is then given by the ratio.

## B Appendix 2 note on more general stoichiometries

Other stoichiometries for mutation/migration reactions can also be considered. In general, a first-order reaction of the form

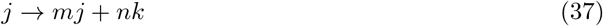

will contribute a term proportional to

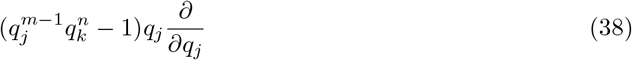

to the Hamiltonian in equation (11). More general mass-action kinetics can also be considered in this framework: if we write 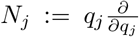 for each reagent *j* and 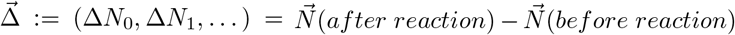 for the overall stoichiometric vector of the reaction, then the most general term for a reaction will be

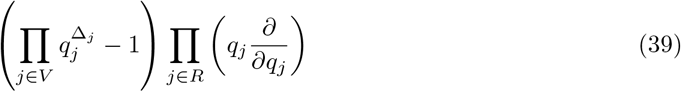

where *R* ⊂ *V* is the set of reagents for the reaction. Note that the various *N*_*j*_ operators commute [27, 32].

## C Appendix 3 projection of level sets of the Fisher information matrix

Our statistical inference framework finds a global minimum of the negative log-likelihood − lnℒ. Futhermore, being able to efficiently evaluate the likelihood function means that we can also estimate the Fisher information matrix from the Hessian *H* of ℒ near the best guess”.

By Wilks’ theorem, we can then generate confidence intervals by studying the level sets of *H*. Our parameter space 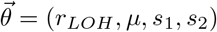 is four-dimensional. To visualise structures in this space, it is natural to ask what two-dimensional projections of these structures look like.

Consider the ellipsoid hypersurface *E* defined by

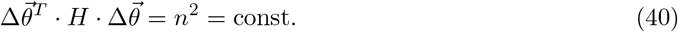

where *n* is an integer that indexes the confidence interval, and

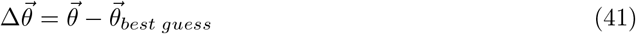

is a vector representing differences from the best guess, or minimum of − ℒ.

Without loss of generality, choose the first two components of the vector 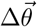 as the *x* − *y* plane onto which we will project an image of the level set defined by (40), and call these 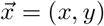. Label the other components of 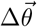 with 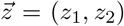. Thus, 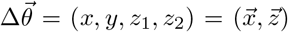. The 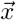 −vector represents coordinates in the image plane *S*, and the 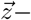 vector represents all other coordinates.

Everywhere on *E*, the surface normal to *E* must lie parallel to the image plane. So for both the *z*_1_ and *z*_2_ directions, we must have the following:

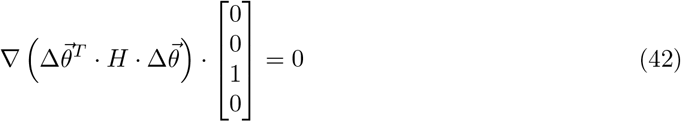

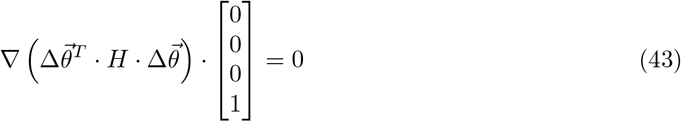

Now,

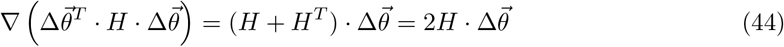

because *H* is symmetric.

Decompose *H* in the following way:

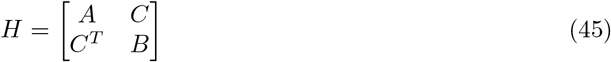

and the two conditions (42) and (43) become

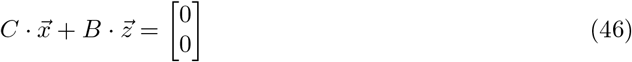

so everywhere on the boundary *E*, 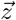 and 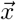 must be linearly related (i.e. coplanar):

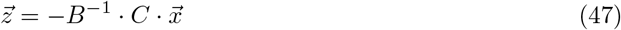

and thus

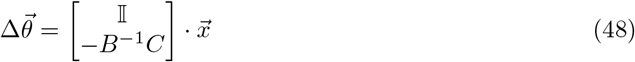

with 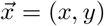; so

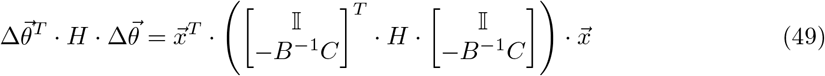

and finally

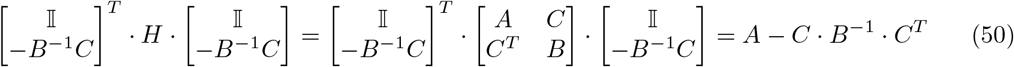

so to visualise the confidence region, the projection of the level set *E* given by (40) can be expressed purely in terms of the image plane coordinates 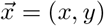 in the following way:

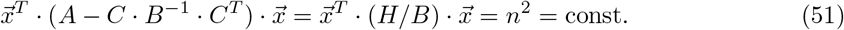

that is, using the Schur complement *H*/*B* [48].

## D Supplemental figures

